# Multi-trophic metacommunity responses to habitat fragmentation in the Brazilian Atlantic Forest

**DOI:** 10.1101/2025.10.03.680099

**Authors:** Tiago Souto Martins Teixeira, Omar Khalilur Rahman, Alyssa R. Cirtwill, Kelly A Speer, David Hemprich-Bennett, Alexis Brown, Susan L. Perkins, Nancy B Simmons, Stephen J Rossiter, Ana Cláudia Delciellos, Marcus V. Vieira, Elizabeth L. Clare

## Abstract

The structure of ecological communities is profoundly altered by anthropogenic disturbance to landscapes. However, most reported impacts rely on the quantification of diversity estimates for single trophic levels or impacts on key species of interest. In this analysis we integrate measures of community structure, comparisons of interaction networks and measures of *β*-diversity across four trophic levels: plants, bats, bat ectoparasites and bacteria within the ectoparasites. Our data show that bat, bat fly, and bacterial communities are significantly nested across forest fragments, with specialist consumers in all groups being found in fewer fragments than generalists. We found substantial *β*-diversity in both species richness and interaction richness across fragments but no decline in interaction redundancy with decreasing fragment size, likely because even intact forest networks had very low redundancy in our dataset. Despite the loss of species and interactions, our data provide support for the conservation value of even the smallest and most disturbed forest fragments where essential seed dispersal services are potentially maintained by *Artibeus*, *Carollia* and *Sturnira* and distinct sets of taxa and interactions are supported. These species may be key in the potential recovery of these habitats, but our data highlight the fragility of these communities which have contracted around these disturbance-tolerant species.

## Introduction

Habitat loss and fragmentation represent global threats to biodiversity, decreasing available habitat while increasing edge effects and isolation among communities (Fahrig, 2003; Ewers & Diidham, 2007; Hagen *et al*., 2012; Püttker *et al*., 2020). In some cases, loss and fragmentation of habitats lead to the formation of discrete local communities within a larger ‘metacommunity’ (Leibold *et al*., 2004), with processes such as resource distribution, niche selection and stochastic drift determining the composition and variability among local communities (Hanski, 1998; de la Sancha *et al*., 2014). Understanding how species respond to habitat loss and fragmentation (Fahrig, 2003; Turner, 1996) remains a key question in the face of large-scale logging and clearing which is disproportionally impacting diverse tropical ecosystems (e.g. Gaveau *et al*., 2014). Most studies have focused on the complex responses of community diversity (e.g. Hertzog *et al*., 2019) and local ecological cascades resulting in extinctions (Portela & Dirzo, 2020) and the loss of genetic diversity (Struebig *et al*., 2011). In a recent mass-scale study of community response to gradients of logging intensity Marsh *et al*. (2025) found that responses were highly variable and non-linear. In their study, some processes like moderate logging had immediate impacts on soil structure, while biodiversity and ecosystem functioning were resilient to logging but affected by conversion to monoculture plantations (Ewers *et al*., 2024b; Marsh *et al*., 2025). Equally important, but more challenging to assess, are the responses of ecological interactions among species (Grass *et al*., 2018) to habitat gradients (Poisot *et al*., 2017).

Fragmentation and habitat loss often disproportionally impact specialists. In Argentinian forests, species loss in smaller fragments led to the contraction of ecological networks with links concentrated around a core of highly-connected species in both bat-plant and host-parasitoid networks (Valladares *et al*., 2012). In the highly-fragmented Atlantic Forest of Brazil, habitat fragmentation led to a decline in specialist pollinator-plant interactions and the disappearance of some pollination syndromes (Girão *et al*., 2007). Habitat loss and climate change are also estimated to disproportionately impact parasites, especially those with complex, multi-host life cycles, as parasites are vulnerable to the compounded effects that habitat loss has on host availability (Cizauskas *et al*., 2017; Carlson *et al*., 2017). The loss of specialists is particularly concerning in areas like the Atlantic Forest, which is a hotspot for biodiversity and endemism (Myers *et al*., 2000) and has suffered extensive habitat loss and fragmentation since the arrival of Europeans *>*500 years ago, accelerating in the 20th century (Costa *et al*., 2017). Only 22.86% of original forest cover remained in 2020, with 97% comprising forest fragments smaller than 50 ha (Vancine *et al*., 2024). Since many of these forest fragments have been established for hundreds of years, the Atlantic Forest is an excellent system for understanding the long-term impacts of habitat modification, with implications for other forests where large-scale habitat fragmentation is ongoing (Gaveau *et al*., 2014).

If endemic species with small ranges are also more likely to be specialist, as suggested by Holt (2010), then a disproportionate loss of specialists in forest fragments is also likely to result in more extinctions among endemic species. However, if generalists are able to persist even in small fragments, this offers the potential for specialists to re-establish themselves from nearby forest fragments. Specialists often depend strongly on generalist interaction partners (Bascompte *et al*., 2006; Aizen *et al*., 2008), even if those generalists are introduced species (Stouffer *et al*., 2014). For example, specialist fruit bats may be able to immigrate to a forest fragment if their food plants are previously seeded there by generalist bats. Though the loss of specialists in a forest fragment impacts fewer interaction partners than the loss of a generalist, it does reduce the amount of functional redundancy in an interaction network. Redundancy is generally believed to promote resilience as species with overlapping sets of interaction partners may ‘compensate’ for each other if one becomes rare (Naeem, 1998; Rosenfeld, 2002; Aizen *et al*., 2012). While the loss of a species with a highly redundant set of interactions may have little impact on the community, losing redundancy makes the network steadily more vulnerable to the loss of additional species (Naeem, 1998; Peralta *et al*., 2014). However, species and interactions lost at one site may be retained elsewhere in the community (Poisot *et al*., 2017), making *β*-diversity within the metacommunity an important contributor to the potential recovery of forest fragments.

Here we investigate how habitat fragmentation has affected a multi-trophic community of plants, bats which disperse their seeds, parasitic bat flies that feed on bats, and microbes living within bat flies that include both putative symbionts and pathogens. We test 1) how the composition and size of each trophic group is related to fragmentation, 2) whether specialists are more strongly affected by fragmentation, and 3) whether the interaction networks in forest fragments share a consistent structure.

## Materials and Methods

### Study area and sample collection

Our samples were collected in the fragmented lowland Atlantic Forest of the Guapi-Macacu River Basin, southeast Brazil, adjacent to the Reserva Ecológica de Guapiaçu (REGUA) (Fig. S1, *Appendix S1*). REGUA comprises a mosaic of pristine and secondary forest and is connected to the Serra dos Órgãos National Park and the Três Picos State Park, forming one of the largest remnants of Atlantic Forest (Ribeiro *et al*., 2009). We sampled at three sites within the continuous forest of the REGUA (hereafter REGUA 1, 2 and 3) and ten surrounding forest fragments that ranged from 20-243ha (hereafter F1-10). The straight-line distance between a forest fragment and its nearest neighbour (not necessarily sampled) ranged from 60-600m and forest fragments represented a mixture of primary and secondary forest confined to tops of hills and steep cliffs (more details in (Delciellos *et al*., 2016; Finotti *et al*., 2012)).

To quantify landscape characteristics, we combined the ESRI base maps in ArcGIS (10.1) with a map of forest cover obtained from Instituto Brasileiro de Geografia e Estatística (IBGE), and a map of forest fragments obtained from SOS Mata Atlântica (www.sosma.org.br). We imported the resulting composite map into Fragstats 3.1 and extracted forest fragment size (hectares), perimiter (m), isolation (distance between nearest neighbour fragments), forest cover within 500m and 1000m, and the proximity index (Gustafson & Parker, 1992) with 500m and 1000m radius buffers. This buffer was selected based on potential daily travel distance for some of the smaller bat taxa. For example, while longer distance movement is known, the mean long axis home range for

*Sturnira lilium* females in Boliva has been estimated at 1323.8m (795.4-1716.3) (Loayza & Loiselle, 2008). Most landscape characteristics were strongly and positively correlated with log fragment size (perimeter, R^2^=0.978; proximity (500m), R^2^=0.979; forest cover (1000m), R^2^=0.919, forest cover (500m), R^2^=0.800; and proximity (1000m), R^2^=0.9786). Isolation was weakly and negatively correlated with log fragment size (R^2^=-0.577); we therefore considered log fragment size, isolation, and the habitat as explanatory variables. Habitat complexity was evaluated by nine habitat variables described in (Delciellos *et al*., 2016). In brief, these were quantified for each forest fragment and sampling area in REGUA and included the presence and absence of watercourses, lianas, *Cecropia* sp., grasses, fallen logs and palm *Astrocaryum aculeatissimum*, and high overstory vertical density, understory horizontal density and tree size ranked as low medium and high. To reduce model dimensionality, we identified principal components axes for habitat complexity using the function ‘prcomp’ in R (R Core Team, 2023). All variables were centered and scaled prior to the principal component analysis.

We captured bats from sunset to midnight at each forest fragment for six nights, from May 2016 to January 2017 using ground-level mist-nets (6, 9 and 12 metres) set along trails, near to streams, and near to flowering or fruiting plants, equating to a net effort of 275,940 m^2^h. We moved nets each night to avoid bats learning their positions. We identified bats following published guides (Emmons & Feer, 1997; Reis *et al*., 2007, 2013) with input from museum taxonomists to confirm identifications. Individual bats were placed in separate clean cotton bags for two to six hours and faecal pellets (guano) produced were stored in ethanol at –20°C. Each bat was searched for ectoparasites for 90 seconds, and bat flies present were collected and preserved in ethanol. Bats were released after the closure of nets in the location of capture. Bats were captured and handled using methods in line with recommendations set out by the American Society of Mammologists (Sikes & Animal Care and Use Committee of the American Society of Mammalogists, 2016) and field work was conducted under Instituto Brasileiro do Meio Ambiente e dos Recursos Naturais Renováveis permit 19037-1.

#### Seed and pollen identification

To identify plants visited by bats we separated guano containing seeds or fruit pulp using a microscope and selected three seeds or one piece of pulp from each. This material was sent to the Canadian Centre for Biodiversity Genomics, University of Guelph, Canada for plant barcoding targeting ITS2 and rbcL regions (Kuzmina *et al*., 2012). We compared the resulting sequences with the reference databases in the Barcode of Life Data System (BOLD) and GenBank following (Lim *et al*., 2018) with the following modifications. We recorded all matches above 95% similarity to a reference and excluded any match not known from the Atlantic Forest of southeast Brazil. For sequences generating a 100% match we retained that identification. For matches *<*100% we separated those sequences generating only one match from those generating multiple matches. If there was only one match or all matches were to members of the same genus, the genus-level ID was retained. If more than one genus was possible between 95 and 100% similarity the sample was excluded due to ambiguity. We compared the identifications retained for both ITS2 and rbcL and if they were the same, we recorded only one bat-plant interaction. If the regions generated different identifications, we assumed each amplified a different target and registered two interactions for that sample. All identifications were then reduced to genus-level, to minimize problems related to different taxonomic resolutions in the network (Hemprich-Bennett *et al*., 2021).

#### Parasite identification

Bat fly identifications are described in (Brown *et al*., 2022; Speer *et al*., 2022). In brief, bat flies were examined under a Leica S9i microscope and first identified using published keys (Theodor, 1967; Wenzel, 1976; Graciolli & de Carvalho, 2001b,a) and by comparison to material from the Field Museum’s collections. Identifications were confirmed by molecular analyses. DNA was extracted using commercially available kits with published modifications (Brown *et al*., 2022; Speer *et al*., 2022). A fragment of the cytochrome oxidase c subunit I gene (COI) was amplified using primers LCO1490 and HCo2198 (Folmer *et al*., 1994), visualised on a 1.5% agarose gel and cleaned using AMPure XP beads. Products were sequenced on the ABI 3730xl DNA Analyzer and the resulting sequences were trimmed to 645bp, checked for quality, and aligned using ClustalW in Geneious v.10.2.4 (Kearse *et al*., 2012). A phylogeny was constructed using RAxML v.8 and a GTR+G model evolution based on AIC scores taken from jModelTest 2.1, with 1000 bootstrap replicates on the CIPRES Science Gateway (Miller *et al*., 2010; Darriba *et al*., 2012). Clades were compared to morphological analysis to confirm bat fly identifications.

#### Microbial identifications

The microbiome communities of bat flies were characterized following (Speer *et al*., 2022). In brief, DNA extractions were performed using commercially available kits developed for low contamination (these extractions were a portion of the ones used to identify parasites above), followed by microbiome sequencing targeting the 16S V4 region using primers 515f and 806r (Caporaso *et al*., 2010). Concentrations were checked using the Qubit 2.0 fluorometer dsDNA HS Assay Kit (Invitrogen) and a Bioanalyzer 2100 DNA High Sensitivity chip (Agilent) to verify quality and amplicon size. Samples were tagged with the Illumina Nextera XT Index Kit v2 (set A, set B, and set C) and multiplexed libraries were sequenced using an Illumina MiSeq v3 Reagent Kit with 2×300bp reads and 18% PhiX spike-in. The reads were de-multiplexed and processed using the QIIME2 pipeline (https://docs.qiime2.org/2018.2/). Chimeras and reads containing PhiX were removed using DADA2, and clustered into amplicon sequence variants (ASVs) (Callahan *et al*., 2016). The GreenGenes Database v13.5 (DeSantis *et al*., 2006) was used for ASV identification. All reads were filtered to remove low resolution data and contaminants using the R (R Core Team, 2023) package decontam (Davis *et al*., 2018). ASVs found in each sample and the taxonomic identification of those ASVs to genus were exported for matrix construction. As parasites were externally washed prior to DNA extraction, we assume that the bacteria are associated with the parasites rather than the bats. This assumption is supported by other research that indicates bat fly and bat skin microbiomes are distinct communities (Speer *et al*., 2024).

### Matrix construction

For each site (three REGUA sites and 10 forest fragment sites), we constructed herbivory, parasitism, and microbial networks that describe interactions between plants and bats, bats and ectoparasitic bat flies, and bat flies and internal bacteria, respectively. Bats and bat flies were identified to species while plants and bacteria were identified to genus. Note that, because bat-plant interactions were detected using plant DNA recovered from bat faeces, we cannot distinguish between frugivory and nectarivory. We therefore combine both types of interactions into a single ‘herbivory’ matrix. As well as these 39 “local” matrices, we also generated three regional metawebs containing all recorded interactions of each type. The weight of each interaction is the number of samples (i.e., bats or ectoparasites) in which the interaction was observed. In addition, we constructed occurrence matrices for each trophic group (bats, ectoparasites, and bacteria). These occurrence matrices included all samples for each group, even those which did not yield interaction data (e.g., bats with no detected ectoparasites or parasites where bacterial DNA processing failed).

### Statistical analyses

#### How does fragmentation affect community size and composition?

We first tested whether the species richness of each consumer trophic group was related to forest fragment properties. We do not model plant genus richness as we lack independent data on plant presence in each site and bats may feed on plants from outside the forest fragment in which they are captured. We expect that the bat community in a forest fragment will be affected directly by habitat changes while the bat fly and bacterial communities will also be affected by changes in the bat community as bat hosts are a major component of their parasites’ habitat (though bat flies are also affected by environmental conditions beyond their bat host (Speer *et al*., 2022)). To test these hypotheses, we modelled bat species richness as a function of habitat characteristics and parasite species richness and microbe genus richness as a function of habitat characteristics and bat richness. Thus, we fit a Poisson regression relating bat richness to log fragment size, isolation, habitat complexity axis 1, and habitat complexity axis 2 and Poisson regressions relating parasite or microbe richness to bat richness and all of the above forest fragment characteristics. All predictors were centered and scaled prior to model fitting; models were fit using the R (R Core Team, 2023) base function ‘glm’. After fitting the full model, we then fit all models including a subset of the predictors and identified the model from this set with the lowest AICc using the function ‘dredge’ from the R (R Core Team, 2023) package MuMIn (Bartoń, 2015). This model indicates the strongest predictor of richness in each trophic group.

Next we tested whether, in general, forest fragments which support few taxa tend to host a subset of the taxa found in fragments with many taxa. That is, we test whether the matrices of species occurrences across fragments are nested. Significant nestedness would be consistent with our hypothesis that some species are more resistant than others to the effects of fragmentation, with the resistant taxa occurring across most fragments and the susceptible taxa occurring in only a subset of the most-suitable fragments. A modular structure, in contrast, would indicate that some fragments are suitable for distinct groups of taxa which are not found in other fragments, while a random structure would indicate haphazard site occupancy and perhaps generally low fragment suitability for these taxa.

We calculated nestedness using the function ‘networklevel’ from the R (R Core Team, 2023) package *bipartite* (Dormann *et al*., 2008). To test whether the nestedness of observed matrices is significant, we compared each observed nestedness to the nestedness of 9999 randomised occurrence matrices. We define the *p*-value for observed nestedness as the proportion of randomised matrices with nestedness greater than the observed value. To create these randomisations, we shuffled the identities of taxa found at each site while preserving the number of taxa found at each site. Thus, we remove any effect of fragmentation on the composition of each community while preserving any effect on community size. We repeated this process separately for plant genera, bat species, ectoparasite species, and bacteria genera.

#### Are specialists most affected by fragmentation?

We expect that, if some taxa are more affected by fragmentation than others, it is relative specialists which will be removed while relative generalists are able to persist in more fragments. For each consumer group (bats, ectoparasites, and bacteria), we calculated the generality of each taxon based on the metaweb. For bats, this is the number of plant genera detected in faecal samples. For ectoparasites, this is the number of bat species on which the parasite was found. For bacteria, we define generality as the number of bat fly species in which the bacterial genus was found.

For each trophic group, we then test whether generality is correlated with the number of fragments (excluding the REGUA sites) in which a taxon was found (site occupancy). To test whether a species’ site occupancy was related to its generality, we first calculated Pearson’s correlation between site occupancy and generality using the R (R Core Team, 2023) base function ‘cor’. Next, we formally tested for significant relationships by fitting a series of Poisson regressions relating occupancy to generality using the R (R Core Team, 2023) base function ‘glm’ and calculating the ANOVA of each model using the R (R Core Team, 2023) base function ‘aov’. We first examined the relationship between occupancy and generality across all trophic groups, and then treated bats, bat flies, and bacteria in isolation and repeated these analyses across all sites (including the three REGUA sites) and the forest fragments only. To supplement these analyses, we also tested whether the mean generality of taxa found in a non-REGUA fragment is related to fragment properties fragment size, isolation, or habitat complexity (*Appendix S2*).

#### Is specialism consistent in different network types?

For bats and bat flies, which were each included in two types of interaction networks, we tested whether generalist consumers in one network were also generalist resources in the other. That is, were bats which consumed many plant genera also hosts to many parasite species and did bat flies which used many bat species as hosts also host many bacterial genera? If so, taxa which were able to establish themselves in a forest fragment by taking advantage of a wider variety of resources might also allow a wider diversity of parasites or bacteria to colonise the fragment. To test this possibility, we first calculated Pearson’s correlation coefficients between bats’ degrees in the herbivory metaweb and parasitism metaweb and between bat flies’ degrees in the parasitism metaweb and microbial metaweb using the R (R Core Team, 2023) base function ‘cor’. We use the metaweb degrees rather than degrees in local observed networks as the metaweb gives a better indication of a species’ true generalism or specialism, unrestricted by a potentially small set of locally-available resources. To test the significance of the correlations, we fit Poisson regressions relating the two sets of degrees (using the R (R Core Team, 2023) base function ‘glm’) and calculated ANOVAs for each (using the R (R Core Team, 2023) base function ‘aov’).

#### How does interaction network structure change between fragments?

Disproportionate loss of specialists could result in reduced redundancy of interaction networks. Another non-exclusive possibility is that changing community composition and environmental context could result in changes to the interaction network over and above changes in species. We investigate evidence for each possibility below.

#### Is loss of redundancy a major effect of fragmentation?

To test for loss of redundancy, we first calculated the mean number of shared partners in each trophic group as an estimate of redundancy. If, on average, more taxa share the same interaction partners, then there is more redundancy in the network. For each trophic group (plants, bats, bat flies, or bacteria) in each network type (herbivory, parasitism, or microbial), we tested whether there was a significant relationship between redundancy and log fragment size, isolation, or habitat complexity (described by the first two PCA axes and their interaction). Mean number of shared partners was calculated using the ‘networklevel’ function in the R (R Core Team, 2023) package *bipartite* (Dormann *et al*., 2008). All models were fit using the base R (R Core Team, 2023) function ‘lm’. In general, redundancy was not related to fragment or site (including REGUA) properties (*Appendix S3*).

#### Did interaction structure change beyond the influence of changing species composition?

To visualise the variability of species and their interactions between sites, we began by calculating Jaccard similarity (J) in consumer species, potential interactions, and observed interactions between each pair of sites (fragments and REGUA sites). Jaccard similarity is simply the number of species common to both fragments divided by the total number of species found at either, or both, sites. Potential interactions were defined as any interaction occurring in the metaweb where the two species involved were observed at the focal site. For herbivory interactions, we assume all plant genera were accessible to bats at any site as plants were not directly sampled. For parasitism and microbial interactions, we considered taxa to be present only if they were observed in this study. Observed interactions were those included in the interaction networks described in the main text.

Note that, because not all samples yielded interaction data, some potential interactions may have occurred but not been observed and our measure of species turnover based on all samples may differ slightly from species turnover calculated from the interaction networks.

To uncover whether changes in the structure of interaction networks was largely due to changes in community composition, we followed (Poisot *et al*., 2012). Using the R (R Core Team, 2023) function ‘betalink’ from the synonymous package (Poisot *et al*., 2012), we calculated the *β*-diversities of species and interactions as well as the proportion of interaction turnover due to rewiring (i.e., not due to species turnover) between each pair of networks of the same type (herbivory, parasitism, or microbial). After calculating the amount of species and interaction turnover and the proportion of interaction turnover due to rewiring, we then tested whether the extent of rewiring was related to the amount of species or interaction turnover. We did this using a set of six linear regressions (two per interaction type) relating the proportion of interaction turnover due to rewiring to the amount of species or interaction turnover and the type of site pair (REGUA-REGUA, REGUA-fragment, or fragment-fragment). We included this term in case turnover or rewiring were generally lower between intact forest sites. All regressions were fit using the R (R Core Team, 2023) base function ‘lm’.

## Results

### Species capture and identification

We recorded 988 captures of 26 bat species (Fig. 1; Table S11, *Appendix S4*). Verified recaptures represented *<*1% of all records. Captures were dominated by the Phyllostomidae with *Carollia perspicillata* the most abundant species with 382 captures. *Artibeus lituratus*, *Artibeus obscurus*, *C. perspicillata* and *Sturnira lilium* were sampled across all sites in contrast to *Chiroderma doriae*, *Micronycteris minuta*, *Tonatia bidens* and *Trachops cirrhosus* captured at only one site each. Four bat species (*Chiroderma doriae*, *Diphylla ecaudata*, *Pygoderma bilabiatum*, and *Tonatia bidens*) were only captured at the REGUA sites and seven bat species (*Anoura caudifer*, *Eptesicus brasiliensis*, *Histiotus velatus*, *Micronycteris minuta*, *Myotis riparius*, *Peropteryx macrotis*, and *Trachops cirrhosus*) were only captured at the fragment sites. A total of 475 seed/pulp samples were processed for DNA barcoding, from which we obtained genus identifications for 354 samples (74.5%). Of the 15 plant genera identified overall, two (*Trichilia* and *Vismia*) were only found in REGUA samples while seven (*Desmodium*, *Musa*, *Myrsine*, *Psidium*, *Sida*, *Syzygium*, and *Terminalia*) were only found in fragment samples.

**Figure 1:**
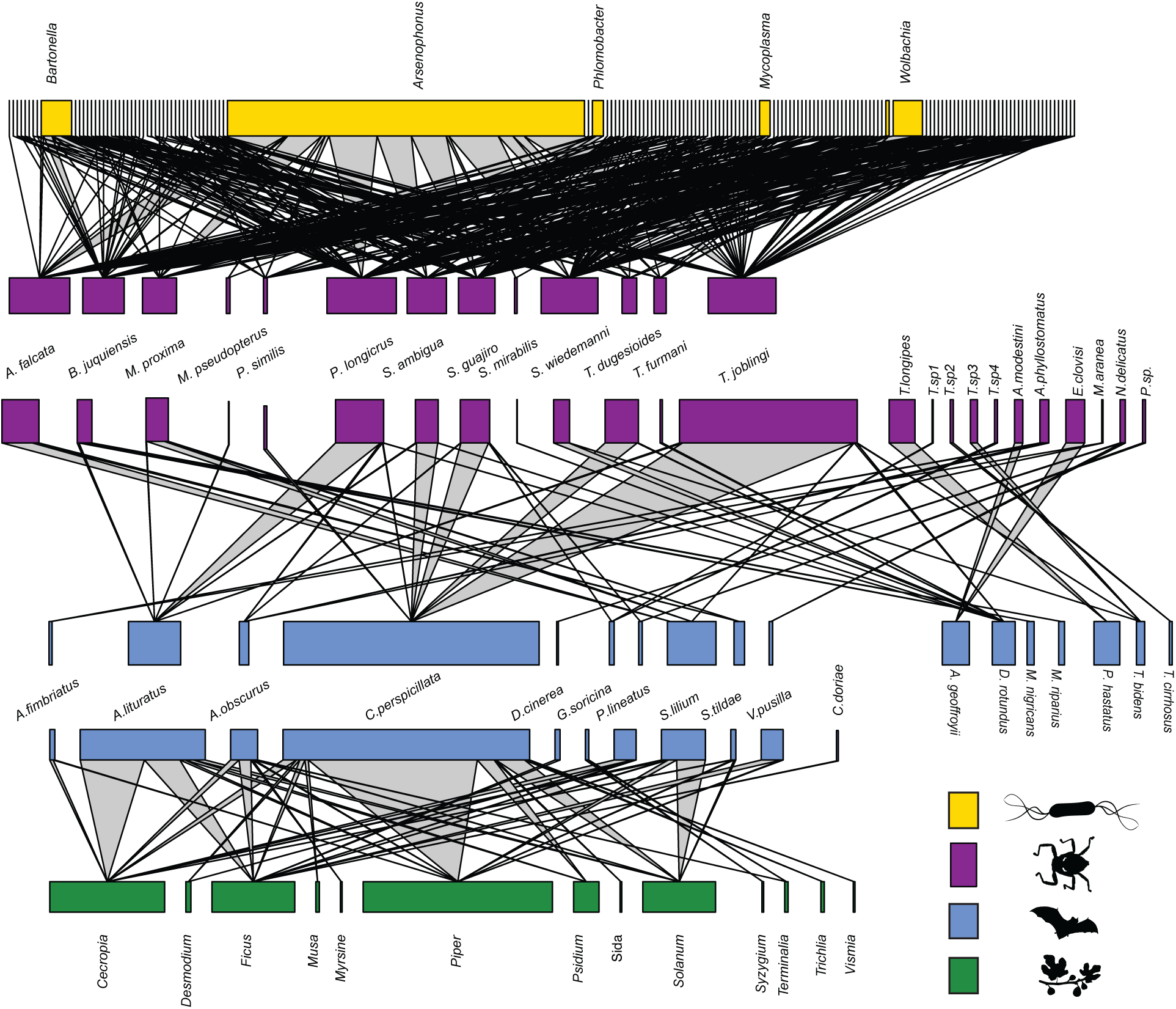
Metawebs describing herbivory (frugivory and/or nectivory), parasitism, and parasite-microbe interactions across 13 sites in the Brazilian Atlantic Forest. This network depicts all recorded interactions among plants (green), bats (blue), dipteran parasites (purple) and their microbes (yellow). Vertical names indicate species found in only one network, angled names indicate species shared between networks. Node width depicts relative presence within a bipartite level.

**Figure 2:**
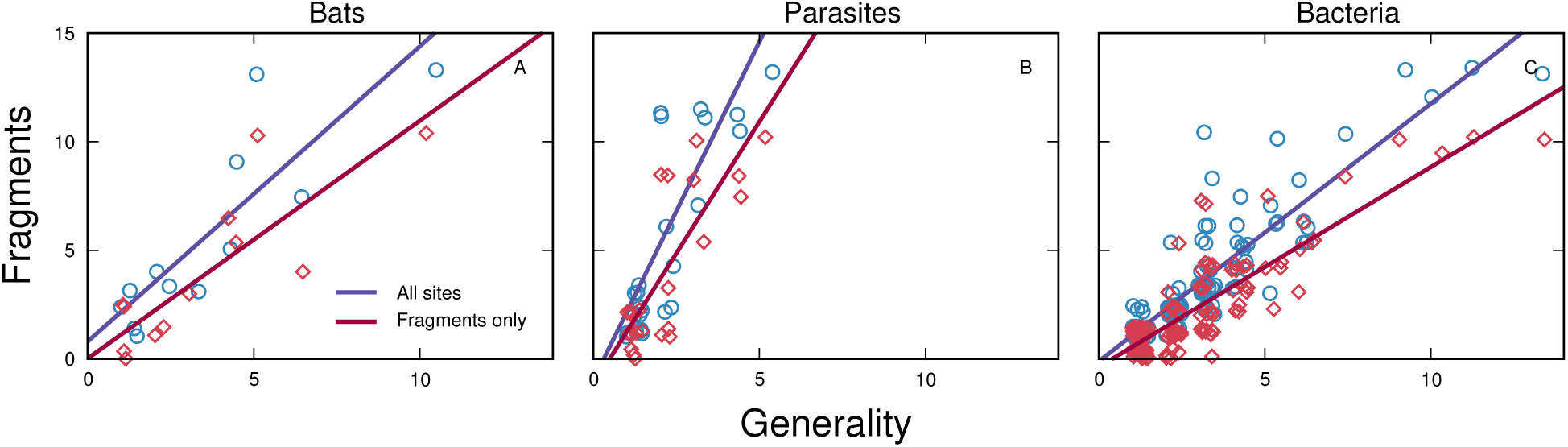
For bats, bat flies, and bacteria, taxa with greater generality tended to occur in more forest fragments. This was true when including all sampling sites (blue, circles) and when excluding the three REGUA sites, which are part of a large continuous forest segment (red, diamonds). Lines represent fits of linear models relating number of sites or fragments in which a taxon was found to its generality.

We collected 842 bat flies (Table S12, *Appendix S4*), of which 436 were found on *C. perspicillata*. Site F7 had the highest incidence of bat fly parasitism (201 bat flies collected). In 29 bat flies, morphological identifications were only possible to genus. Subsequent molecular analysis and comparison to reference material in DNA databases supports the assignment of four of these to species, the remaining 24 to four MOTUs based on reciprocally monophyletic clusters in our phylogeny which we refer to as *Trichobius* sp.1, *Trichobius* sp.2, etc. and treated as presumptive species. The most abundant bat fly across all fragments was *Trichobius joblingi* which accounted for 303 of the 842 bat flies. Three bat flies (*Metalasmus pseudopterus*, *Strebla mirabilis*, and *Trichobius* sp. 3) were found only in the REGUA samples while six (*Paratrichobius* sp., *Trichobius angulatus*, *Trichobius longipes*, *Trichobius* sp.1, *Trichobius* sp.2, and *Trichobius* sp.4) were found only in the fragment samples.

We identified 1101 bacterial ASVs from 188 individual bat flies, which were identified as belonging to 173 bacterial genera. ASVs in the genus *Arsenophonus* were the most common in every sample, except specimens of the bat fly genus *Basilia*. Other prevalent bacteria with high relative abundance were *Wolbachia*, *Bartonella*, *Mycoplasma*, candidatus *Phlomobacter*, *Rickettsia*, and *Finegoldia*. Most ASVs were either extremely low in relative read abundance (*<*0.01) and prevalence or cannot be identified to genus based on existing databases. Additional details on patterns of microbial diversity are reported (Speer *et al*., 2022). In total, we recorded 124,832 interactions between bacterial genera and ectoparasites. There were 36 bacterial genera observed only in the REGUA samples and 57 genera recorded only from fragment samples.

### Fragmentation affected community composition but not size

Prior to testing whether habitat fragmentation affected species richness, we used a PCA to condense habitat complexity into two main axes. The first axis explained 35.7% of the variance and was most strongly positively associated with understory density, canopy density, and watercourses, and negatively associated with tree size (Figure S2, *Appendix S1*). The second axis explained a further 27.9% of the variance and was most strongly positively associated with the abundance of *Astrocaryum aculeatissimum* and negatively associated with the abundance of *Cecropia sp*. For comparison, the third axis explained 14.4% of the variance, and the remaining six axes explained less than 10% of the variance.

The best-fit model for bat species richness included only an intercept while parasite species richness was best predicted by bat richness (Table S13, *Appendix S5*). Bacteria genus richness was best predicted by a model including isolation and log fragment size. No other model was within *δAIC_c_<*2 of the best-fit model. There were more bacteria genera in larger and more isolated fragments.

The plant, bat, parasite, and bacteria occurrence matrices were all significantly nested compared to randomised occurrence matrices with the same taxon richness per site (Table 1). That is, the taxa found in lower-richness fragments tended to be subsets of the taxa found in higher-richness fragments.

**Table 1:**
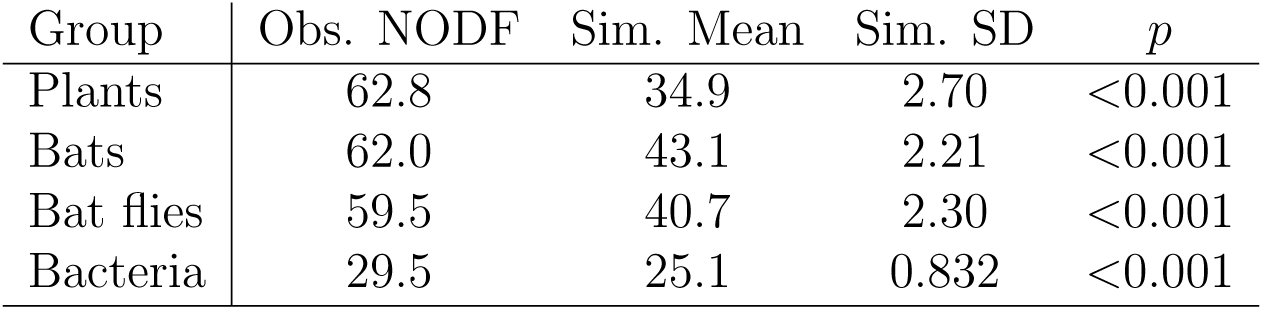
Observed nestedness (NODF) of site-occupancy matrices for each taxonomic group alongside summary statistics of 1000 randomisations of each observed matrix. *p*-values for significance of observed nestedness were defined as the percentage of simulated matrices with equal or greater nestedness to the observed.

| Group | Obs. NODF | Sim. Mean | Sim. SD | $p$ |
| --- | --- | --- | --- | --- |
| Plants | 62.8 | 34.9 | 2.70 | <0.001 |
| Bats | 62.0 | 43.1 | 2.21 | <0.001 |
| Bat flies | 59.5 | 40.7 | 2.30 | <0.001 |
| Bacteria | 29.5 | 25.1 | 0.832 | <0.001 |

### Specialists appear at fewer sites

At all levels, generalist taxa were less affected by forest fragmentation. Plants found in the faeces from a large number of bat taxa also tended to occur in more sites (*β*=0.265, *p<*0.001). Bats which consumed larger numbers of plant genera tended to occur in more sites (*β*=0.136, *p<*0.001). Notably, the four bats which were found at all sites (*Artibeus lituratus*, *A. obscurus*, *Carollia perspicillata* and *Sturnira lilium*) were also the most generalist (consuming 10, 6, 5, and 4 plant genera respectively). Parasites found on a larger numbers of bat host species also tended to occur in more sites (*β*=0.485, *p<*0.001). Bacteria found in larger numbers of bat flies tended to occur in more sites (*β*=0.229, *p<*0.001). Despite these taxon-level trends, the mean generality at a fragment was not strongly related to forest fragment properties (*Appendix S2*).

#### Generalist consumers tend to be generalist resources

Bats’ degrees as herbivores were strongly and positively correlated with their degrees as hosts for bat flies (*R*^2^=0.819, *F*_1,8_=16.3, *p*=0.004; Fig. 3). Likewise, bat flies’ degrees as parasites were strongly and positively correlated with their degrees as hosts for microbes (*R*^2^=0.750, *F*_1,11_=14.2, *p*=0.003). This suggests that taxa which are able to colonise to a forest fragment by utilising a wide variety of food sources or hosts may also facilitate colonisation by a wide variety of parasites or microbes.

**Figure 3:**
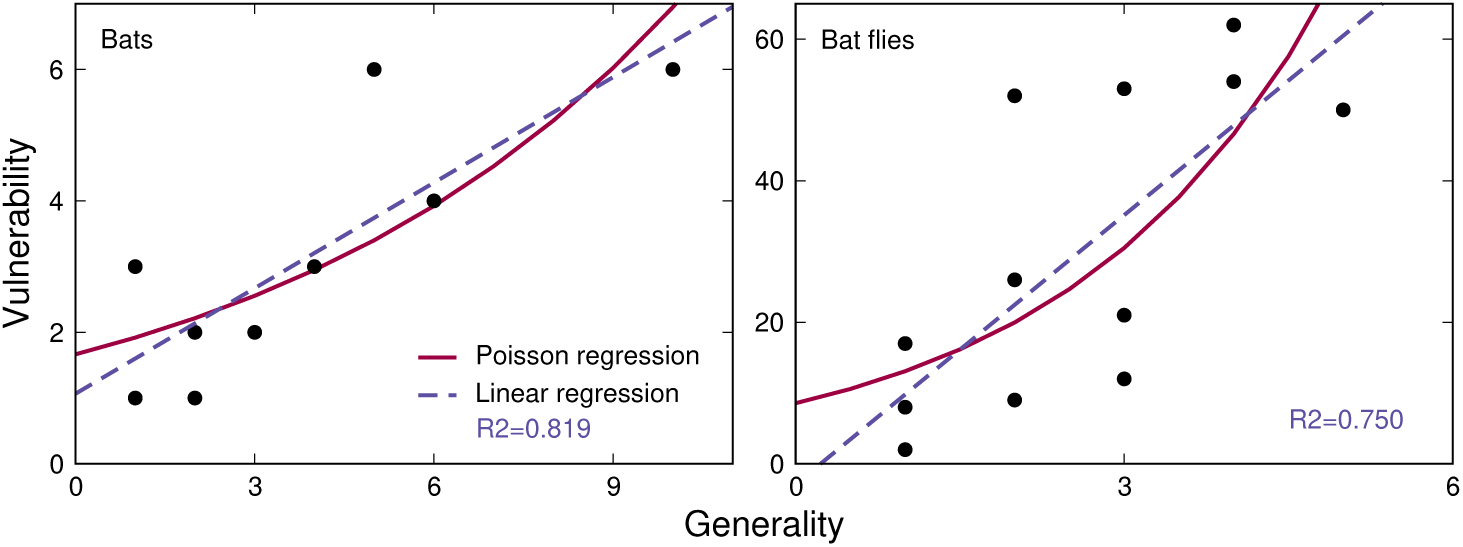
Bats’ and bat flies’ degrees as consumers and as resources were strongly and positively correlated. A generalist bat species tends to support many parasite species and a generalist bat fly tends to contain a wide diversity of microbes. *R*^2^ values given refer to the linear regressions indicated by blue, dashed lines. As degrees are counts of interaction partners, we also fit Poisson regressions (reg, solid lines). Significance did not change depending on the error distribution used.

### Interaction networks showed low redundancy but high turnover

Plants, bats, bat flies, and bacteria were all generally less redundant than expected in their sets of interaction partners (*Appendix S3*). The only exceptions were bacteria at sites F2, F3, and F6, which had significantly more shared bat fly hosts than expected, and bat flies at site F2, which had significantly more shared bacterial genera than expected. Perhaps because of this generally low redundancy, there was no relationship between redundancy and site properties for most trophic groups and network types (*Appendix S3*). The one exception was the parasitism networks where, if the REGUA sites are included in the analysis, bat flies were less redundant in their bat hosts in larger fragments (*β*=-0.051, *p*=0.024) while the redundancy of parasite communities across bats was related to habitat complexity (Table S2, *Appendix S3*).

Most of the observed interaction matrices were significantly more nested than the mean nestedness of 1000 randomisations of the network (*Appendix S3*). This was true for both binary and weighted nestedness. The exceptions to this trend included parasitism and microbial networks from both REGUA and fragment sites. However, nestedness was not related to fragment size, isolation, or habitat complexity unless the three REGUA sites were included, when nestedness in the parasitism networks was related to habitat complexity (*Appendix S3*).

Overall, Jaccard similarity in species turnover between sites was higher for bats and parasites than for microbes (Fig. 4). There was no clear difference in similarity between REGUA sites and forest fragments. Similarity in potential interactions between sites was much higher for herbivory interactions than parasitism or microbial interactions because we assume that bats can always access any plant recorded in our dataset. This assumption is likely false but, without data on the plant species found in each fragment, we have no reasonable alternative. Similarity was also higher in potential parasitism interactions than potential microbial interactions, likely because of the higher turnover in the microbial community. Realised interactions were less similar between sites than potential interactions in all cases, but most notably for herbivory interactions due to the assumption of constant plant availability mentioned above.

**Figure 4:**
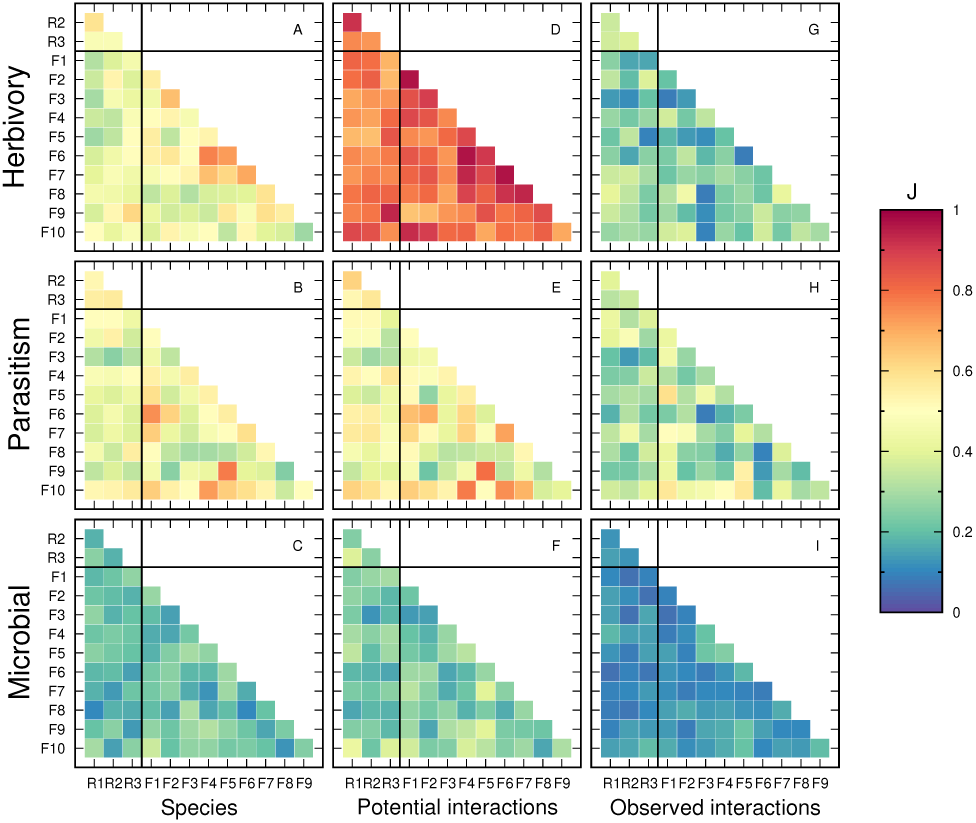
Jaccard similarity between consumer species composition (A, B, C), potential interactions (D, E, F), and realised interactions (C, H, I). In each case, the taxon indicated on the row label is the consumer in the interaction (e.g., bats are consumers in herbivory interactions). Note that, when calculating potential herbivory interactions, we assume that bats can access all plant genera from all forest fragments. This assumption greatly increases the similarity of potential interactions across sites. Species and potential interaction similarity is based on all samples, including those which did not yield interaction data. Horizontal and vertical lines in each panel distinguish between REGUA (top and left, labels beginning with ‘R’) and fragment (lower and right, labels beginning with ‘F’) sites.

In all three network types, the proportion of interaction turnover due to rewiring decreased significantly with increasing species turnover, though there was substantial variation around this trend (Table S14, *Appendix S6*; Figure 5). For herbivory and parasitism networks, but not microbial networks, the proportion of interaction turnover due to rewiring tended to increase with increasing interaction turnover.

**Figure 5:**
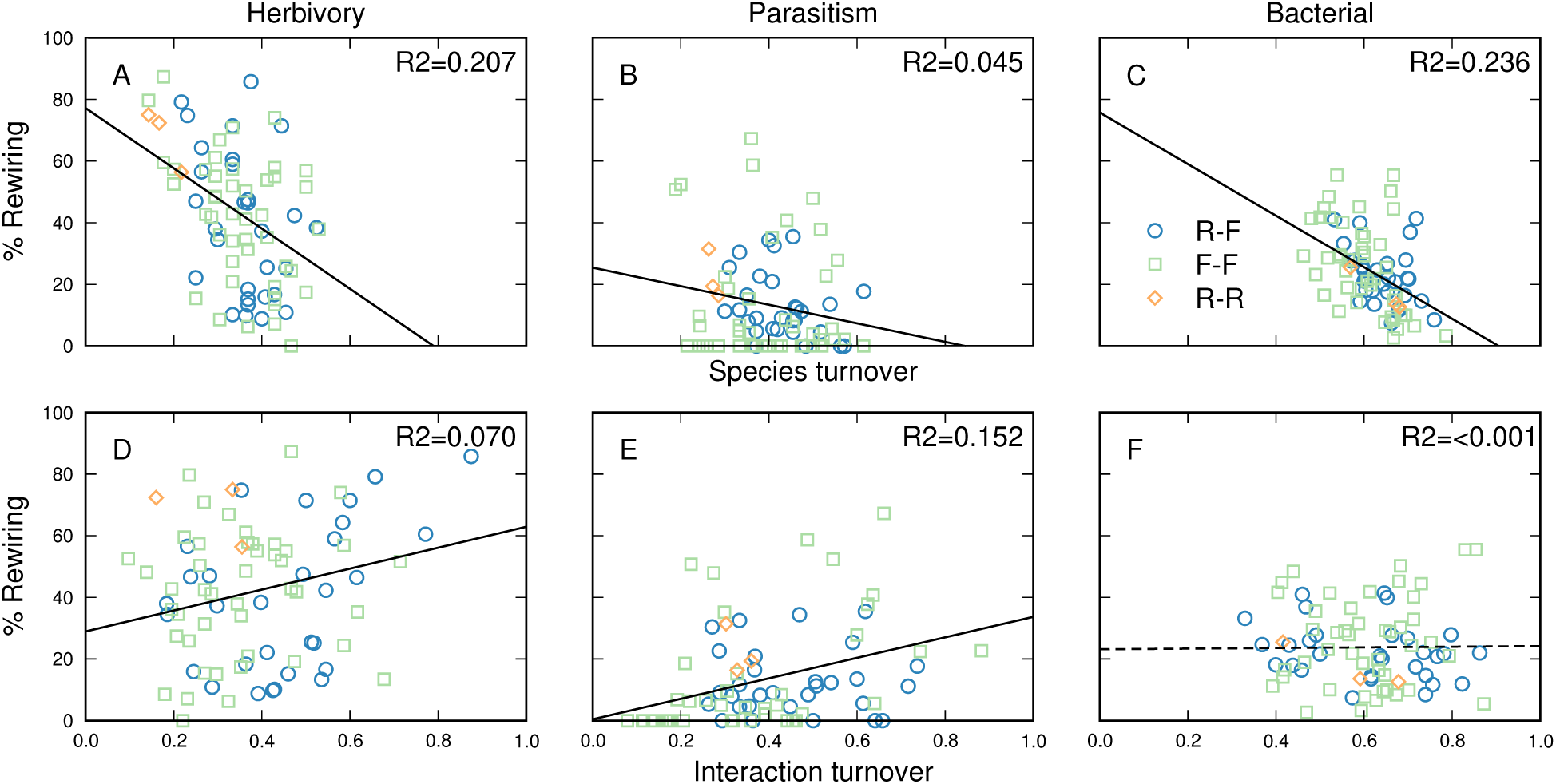
All three network types showed substantial species turnover (top) and even more interaction turnover (bottom). There was no clear difference in the amount of turnover between pairs of fragments (F-F, green squares), pairs of REGUA control sites (R-R, orange diamonds), or control-fragment pairs (R-F, blue circles). The proportion of interaction turnover due to rewiring decreased with increasing species turnover in all network types and increased with increasing interaction turnover for herbivory and parasitism networks.

## Discussion

Our objective was to quantify the impact of habitat fragmentation across a multi-trophic community of bats, the seeds they disperse, their parasites and the microbiome of the parasites. Our analysis finds considerable contraction in community composition and loss of interactions in forest fragments, but also highlights the resilience of key disturbance-tolerant species which provide important forest regenerative services even in the smallest fragments. This provides strong support for the conservation of even the smallest fragments but emphasizes the fragility of these communities.

### Fragmentation changes community composition but rarely size

We did not record any significant changes in bat species richness with fragment size, isolation, or habitat complexity, however the bat community was significantly nested across forest fragments. This means that bats present in depauperate fragments tend to be a consistent subset of the bats present in more species-rich fragments. In this case, the core of widespread bat species (*Carollia perspicillata*, *Artibeus lituratus*, *A. obscurus*, and *Sturnira lilium*) are all common neotropical phyllostomids, abundant in the Atlantic Forest (Muylaert *et al*., 2017) and very tolerant to disturbance (Emmons & Feer, 1997; Reis *et al*., 2007). The fact that they are present even in the most species-poor fragments highlights their importance in maintaining critical ecological functions that are key to the potential recovery of fragmented tropical forests. The *Carollia*, *Artibeus* and *Sturnira* are well-known for their role in primary seed dispersal in neotropical forests, moving seeds between foraging areas in different habitats (Kunz *et al*., 2011). Their persistence in even the most degraded forest sites is particularly important, given that these bat species were highly generalist consumers of plant genera and highly generalist hosts of bat flies, providing support for trophic levels above and below them. However, the very small core of bat species also means that the seed dispersal services they provide is likely vulnerable to any decline in the abundance of these bats due to disease, anthropogenic impacts, climate change, or habitat degradation.

The bat fly and endobacterial communities were also significantly nested across forest fragments. The number of bat fly species present increased with increasing bat species richness, reflecting the requirements of specific hosts for some parasites. As with bats, generalist bat flies tended to occur in more sites than specialists. Even the most generalist bat flies tended to parasitize only a few bat species (up to 5) compared to the diet breadth of generalist bats which may consume up to 10 or more plant genera. Nearly half of the bat flies recorded (12/25) parasitised at least one one of the four core bat species found in every forest fragment. These included both specialists such as *Paraeuctenodes similis* (found only on *Carollia perspicillata*) and *Metalasmus pseudopterus* (found only on *Artibeus lituratus*) and generalists such as *Speiseria ambigua* (found on *C. perspicillata, A. liturbatus, and Desmodus rotundus*), *Strebla guajiro* (found on *C. perspicillata, A. obscurus, Glossophaga soricina, and Platyrrhinus lineatus*), and *Trichobius joblingi* (found on *C. perspicillata, S. lilium, A. liturbatus, Tonatia bidens, and Desmodus rotundus*). We note that *Carollia perspicillata* hosted over half of all observed bat fly individuals. As this species is known to move 1-2km between habitats for foraging (Kunz *et al*., 2011), it is likely crucial for dispersal of bat flies between habitat patches. As viruses and endoparasites are known to affect the outcome of competition in several systems (e.g., squirrels in Tompkins *et al*. (2002), birds in Tompkins *et al*. (2001), and cervids in Bender *et al*. (2005)), it is also possible that bat flies moderate competition between *Carollia* and other, more specialist, bat species (though our data do not allow us to test this possibility).

Bacterial richness of bat flies was not related to bat host richness but was higher in larger and more-isolated forest fragments. Because bat fly microbiome can be both vertically inherited and or horizontally acquired (see Speer *et al*. (2022) and references therein), bat flies on hosts in more isolated fragments may acquire a wider variety of bacteria from sources in the agricultural matrix around the forest fragment if bats and their parasites pass through these areas on feeding flights. In a separate analysis of the microbial networks using the same data, Speer *et al*. (2022) found that parasite species was a significant determinant of microbial community composition, suggesting that the identities of bat flies found in large and isolated forest fragments may also strongly shape bacterial richness.

In their analysis, (Speer *et al*., 2022) found that bacterial network topology also varied with fragment size. Specifically, bacterial association networks in smaller fragments tended to be less modular, with higher average centrality of bacteria in them, with less-abundant taxa being the most central (Speer *et al*., 2022). Combined with our finding of extremely high turnover in the bacterial community between fragments, the connective role of rare bacteria emphasises the importance of conserving both large and small habitat fragments to protect bacterial diversity. Bacteria contribute a substantial proportion of Earth’s biodiversity (Locey & Lennon, 2016) and are vital for many ecosystem functions (Averill *et al*., 2022; Blaser, 2017), benefit wildlife populations (Trevelline *et al*., 2019), and are necessary associates to ensure the survival and reproduction of their hosts, specifically those that occupy narrow dietary niches (Feldhaar & Gross, 2009). Bacteria are also vulnerable to co-extinctions along with their hosts (Weinbauer & Rassoulzadegan, 2007) and independent responses to threats such as environmental change (Averill *et al*., 2022). In this context and considering the nested structure of bacterial occurrence, preserving large and isolated forest fragments may be worthwhile to safeguard the greater bacterial diversity within them as well as for the habitat offered to plants and animals (Averill *et al*., 2022).

### Extensive turnover in interactions but little change in redundancy

All three network types we considered were characterised by high *β*-diversity in both species and interactions between fragments with low redundancy in species’ interaction partners. Though we expected a disproportionate loss of specialists from forest fragments to lead to reduced redundancy, instead it appears that much of the redundancy in these networks has *already* been lost. Thus, we do not observe significant change in redundancy with fragment size, isolation, or habitat complexity.

Species turnover was high among bats, bat flies, and bacteria, as has previously been shown for frugivorous birds in the Atlantic Forest (Emer *et al*., 2018). We did not quantify species turnover in the plant community as we do not have independent data on which plants were present in which fragment and bats may consume fruit from outside sources, but it is reasonable to expect some variability in the plants present at each site as well.

Turnover was especially high among bacterial genera which were also, on average, observed at fewer sites than bats or bat flies (mean = 2.65 for bacteria, 5.33 for bats, and 4.88 for bat flies).

There was even greater turnover in interactions, indicating changes in interactions due to both changes in community composition and rewiring of interactions between species that co-occur at multiple sites (Poisot *et al*., 2012). Rewiring can occur due to differences in local abundances or species traits, or due to external influences such as environmental conditions and the presence of other species not involved in the interaction (Poisot *et al*., 2015, 2017; Clegg *et al*., 2018). The proportion of interaction turnover due to rewiring was especially high in the herbivory networks, suggesting that bats are relatively flexible in the plants they consume, potentially facilitating re-colonisation of additional forest fragments if dispersal is facilitated. The ecological flexibility of bats is increasingly being recognized both within and between traditional “guild” designations (Clare & Oelbaum, 2024). The generally lower proportions of interaction turnover due to rewiring in the parasitism and microbial networks indicates that bat flies and their endobacteria are somewhat more restricted in the sets of partners that they can use. As many bat fly species are host-specific (Dick, 2007), it is not surprising that they would have stable interactions with a given bat host species. Additionally, the most dominant member of the bacteria detected, *Arsenophonus*, is a known obligate bacterial endosymbiont of bat flies (Nováková *et al*., 2009; Trowbridge *et al*., 2006; Hosokawa *et al*., 2012). This genus likely provisions B-vitamins that are missing from blood-meal, as has been indicated in other blood-feeding arthropod-bacteria interactions (Santos-Garcia *et al*., 2018; Morse *et al*., 2013; Hosokawa *et al*., 2012). Alternatively, the amount of interaction turnover due to rewiring in the bacterial networks may be due to the very high turnover in the low-abundance members of bacteria communities between sites.

Overall, our results indicate that each forest fragment contains a somewhat unique set of species and interactions, with even the three ‘intact forest’ sites within the REGUA showing substantial amounts of *β*-diversity and interaction rewiring, with the metacommunity structures consistent with random species loss in small fragments (*Appendix S7*). This is consistent with analyses of bird-mediated seed dispersal networks in the Atlantic Forest, where each forest fragment contains unique assemblages of birds and bird-plant interactions (Emer *et al*., 2018) and interaction networks tend to be highly modular, without clear structural trends over environmental gradients (Menezes Pinto *et al*., 2021). In particular, interactions between large bird species were locally lost in fragments *<*10,000 ha and interactions of contrasting sizes (e.g. large birds carrying small seeds) were lost in the smallest forest fragments (Emer *et al*., 2020). This may be an interesting contrast with bats where *Artibeus*, one of the larger-bodied genera, persisted in these small fragments, and they may preferentially carry large fruits.

The Atlantic Forest is a unique mosaic of large land masses intermixed with well-established forest fragments forming a stepping stone system which may aid in the dispersal of species across the landscape. It is concerning then that in both the bird example (Emer *et al*., 2020) and these bat communities, forest fragments showed distinct contraction to particularly disturbance-tolerant species (Emer *et al*., 2018). This downscaling of networks in smaller forest fragments and, in particular, the lack of redundancy in interactions, makes these networks vulnerable to the loss of ecological functions in the long run if interactions continue to be lost from forest fragments.

## Conclusions

This study joins a growing body of research showing that both communities of species and the interactions among them can be highly variable over space (Poisot *et al*., 2015; Emer *et al*., 2020; Pellissier *et al*., 2018; Carstensen *et al*., 2014) or time (Lopez *et al*., 2017; Cirtwill *et al*., 2023; CaraDonna & Waser, 2020). In this context, it is difficult to predict the set of interactions in a fragment either based on a regional metaweb or on the set of species locally present. This may make restoration efforts more difficult if a particular species or interaction is a target. On the other hand, our results suggest that even small forest fragments are worth preserving due to the unique species and interactions within them and echos recent meta-analyses calling for the conservation of even partially degraded habitats (Ewers *et al*., 2024a). Moreover, the extensive rewiring of interactions between fragments and greater persistence of generalist bats and their interaction partners suggests that, if additional species are able to disperse to new forest fragments, they will find interaction partners and be able to persist. Thus, forest fragments may still be able to return to a more natural state even after centuries of fragmentation.

## Acknowledgements

This project could not have been completed without field assistance from Victor Miraglia, Alexandre Polletini, Mario Reis, Roberto Leonan, Liliane Alves, laboratory support from REGUA staff, Marcelo Weksler, Museu Nacional/UFRJ and analytical support form Rob Knell. We particularly appreciate the time and expertise of staff at the Museu Nacional (UFRJ) João Alves de Oliveira and Marcelo Weksler and taxonomist Roberto L.M. Novaes. The research was enabled by funding from the Natural Sciences and Engineering Research Council of Canada through the Discovery Grants Program (RGPIN-2021-03611), support from the Government of Canada’s New Frontiers in Research Fund (NFRFT-2020-0073) and by support from Genome Canada and Ontario Genomics to BIOSCAN-Canada through the Large Scale Applied Research Program to ELC and by funding from Queen Mary University of London to ELC. Lab work funding for KAS and AMB was provided by the American Museum of Natural History’s Richard Gilder Graduate School Student Research Fellowship and Sackler Institute of Comparative Genomics Faculty Funding, KAS was also funded by the Smithsonian Institution Biodiversity Genomics Postdoctoral Fellowship, TSMT was funded by a PhD fellowship from Coordenação de Aperfeiçoamento de Pessoal de Nível Superior (CAPES) through the “Science Without Borders” programme (Brazil). OKR was funded by a scholarship from Diversified Villa Sdn Bhd. ACD had a postdoctoral scholarship from CAPES – Finance Code 001 (PNPD-PPGEE/UERJ, project number 1631/2018) and has a scholarship from Universidade do Estado do Rio de Janeiro/FAPERJ (PAPD Docência). MVV was supported by grants from CNPq (312465/2019-0) and FAPERJ (203.045/2017).

## Data accessibility

Parasite and microbial data are available from Speer *et al*. (2022) and (Brown *et al*., 2022). Plant data are available on the BOLD data system and will be made public on manuscript acceptance.

