## Appendix for "Multi-trophic metacommunity responses to habitat fragmentation in the Brazilian Atlantic Forest"

1 School of Biological and Chemical Sciences, Queen Mary University of London, London, UK
2 Carex EcoLogics, Bracebridge, Canada
3 Richard Gilder Graduate School, American Museum of Natural History, New York, NY, USA
4 Department of Biological Sciences, Northern Arizona University, Flagstaff, AZ, USA
5 Pathogen and Microbiome Institute, Northern Arizona University, Flagstaff, AZ, USA
6 Department of Biology, University of Oxford, Oxford, UK
7 Ecology, Evolution, and Environmental Biology, Columbia University, New York, NY, USA
8 Department of Ecology and Evolution, Stony Brook University, Stony Brook, NY, USA
9 Institute for Comparative Genomics, American Museum of Natural History, New York, NY, USA
10 Department of Mammalogy, American Museum of Natural History, New York, NY, USA
11 Departamento de Ecologia, Instituto de Biologia Roberto Alcântara Gomes, Universidade do Estado do Rio de Janeiro, Brazil
12 Departamento de Ecologia, Instituto de Biologia, Universidade Federal do Rio de Janeiro, Brazil
13 Department of Biology, York University, Toronto Ontario, Canada
* Should be considered shared first authors

### Table of contents


1. Site and habitat structure: supplemental figures

2. Mean generality and fragment properties: supplemental methods and results

3. Redundancy and fragment properties: supplemental methods and results

4. Summary of samples: supplemental results

5. Species richness and fragment properties: supplemental results

6. Species and interaction turnover: supplemental results

7. Elements of metacommunity structure (EMS): supplemental methods and results

### Appendix 1: Site and habitat structure

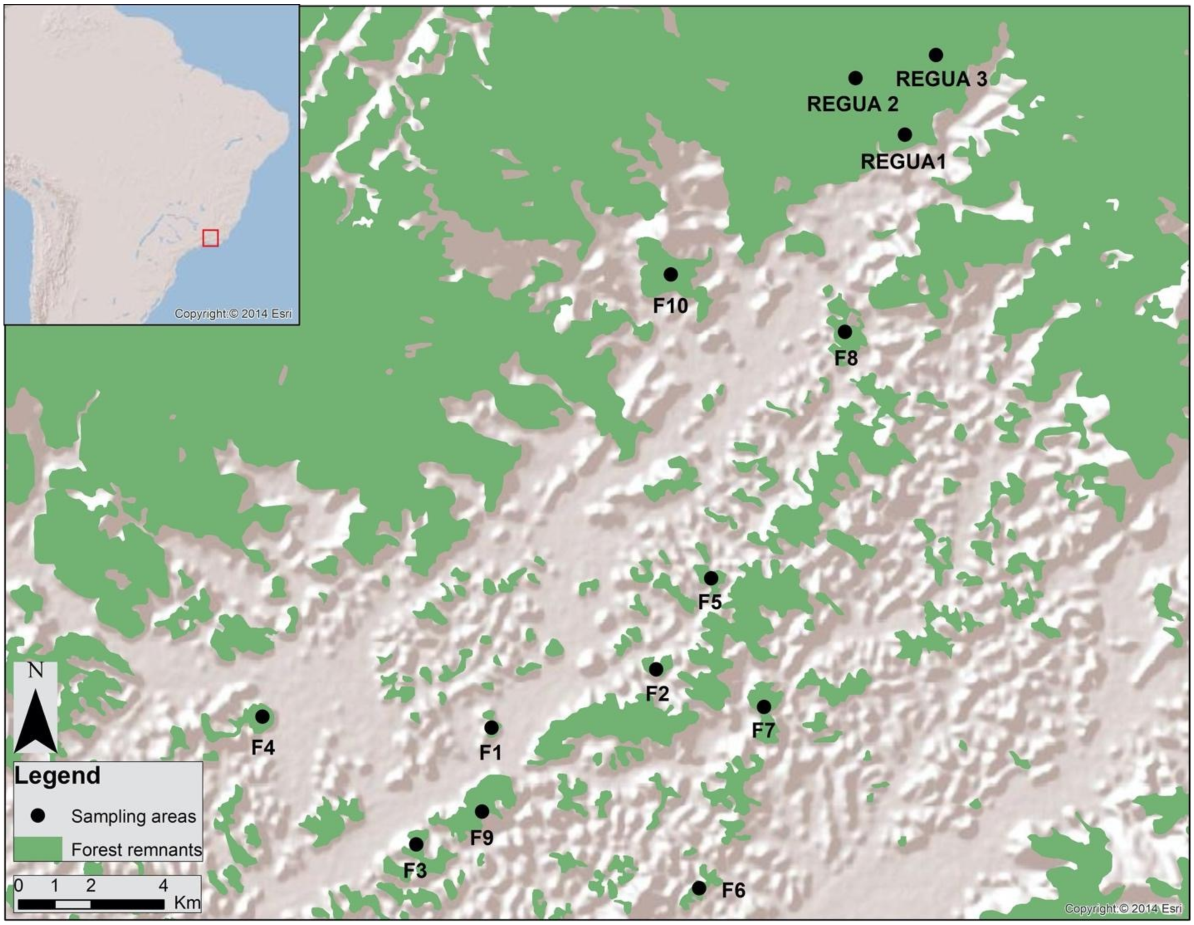


**Figure S1:** Location of sampling sites in a fragmented landscape of lowland Atlantic Forest in the Guapiaçú River Basin of southeast Brazil. We collected samples at three sites the Reserva Ecológica de Guapiaçú (REGUA) and in ten fragments in the surrounding landscape.

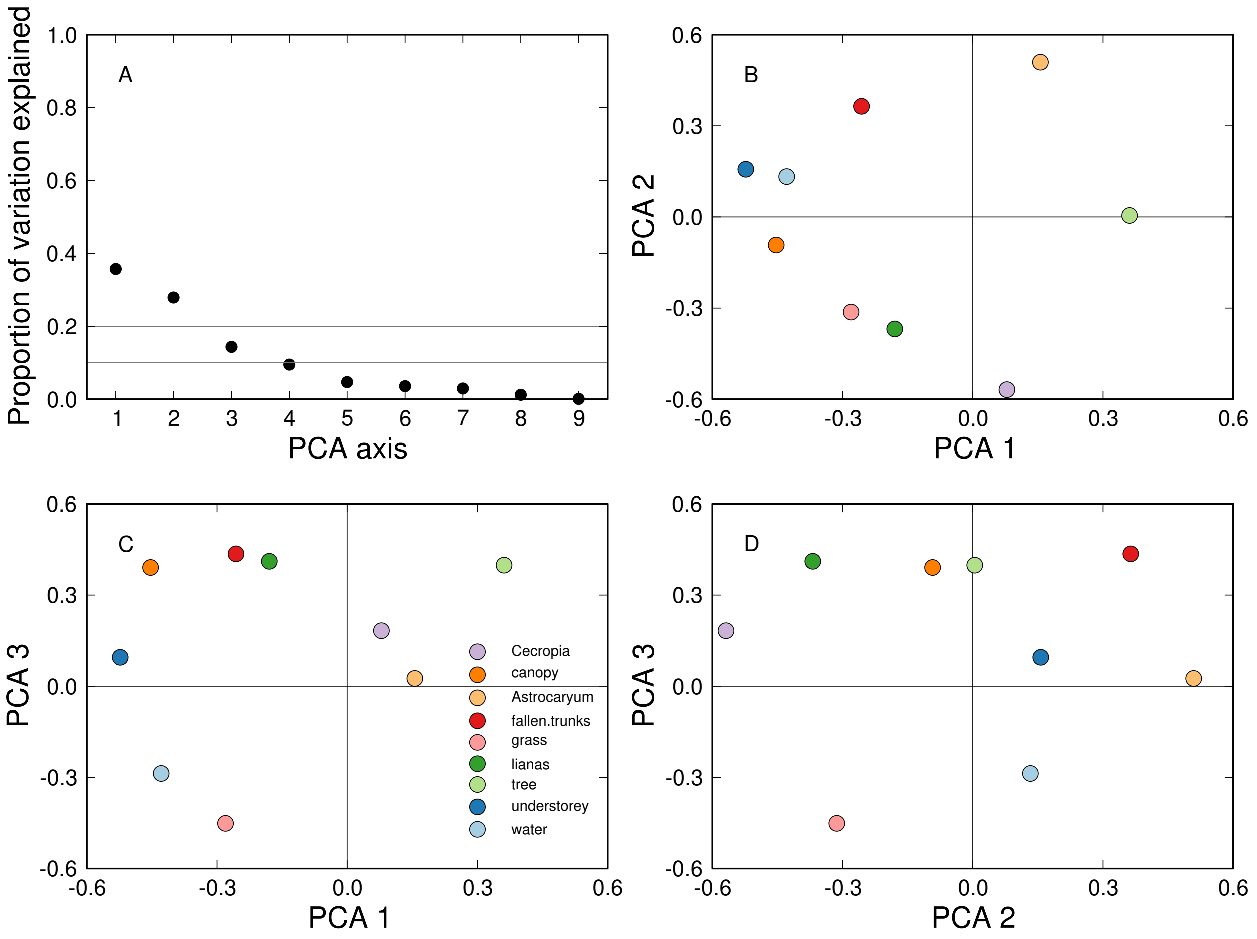


**Figure S2: A)** Proportion of variance explained by all principal components axes for habitat structure, with **B-D)** distribution of measures of habitat structure along the first three PCA axes.

### Appendix 2: Mean generality and fragment properties

#### Methods

As well as testing whether generalists tended to occur at more sites than specialists (*Main Text*), we tested whether the mean generality across all taxa in a group (bats, bat flies, or bacteria) was related to the size, isolation, or habitat structure of a fragment. We first considered all taxonomic groups together and fit a full model relating mean generality to the log area of a fragment, its isolation, the first two habitat PCA axes, and all interactions between them using the R (R Core Team, 2023) base function 'lm' We then identified the best-fitting model (based on AIC) using the R (R Core Team, 2023) function 'dredge' from the package *MuMIn* (Bartoń, 2015). When considering each trophic group separately, the sample size of ten sites did not permit this approach. We therefore fit separate models relating mean generality to log area or isolation or the first two habitat axes and their interaction, all using the R (R Core Team, 2023) base function 'lm'.

#### Results

Mean generality was not related to fragment size, isolation, or habitat complexity. The best-fit model for mean degree included only an intercept and the next-best model, which included an effect of log area, did not show a significant relationship to mean generality ($F_{1,28}$=0.236, $p$=0.631, $R^2$=0.008).

For bats, there was no relationship between mean generality at a fragment and fragment

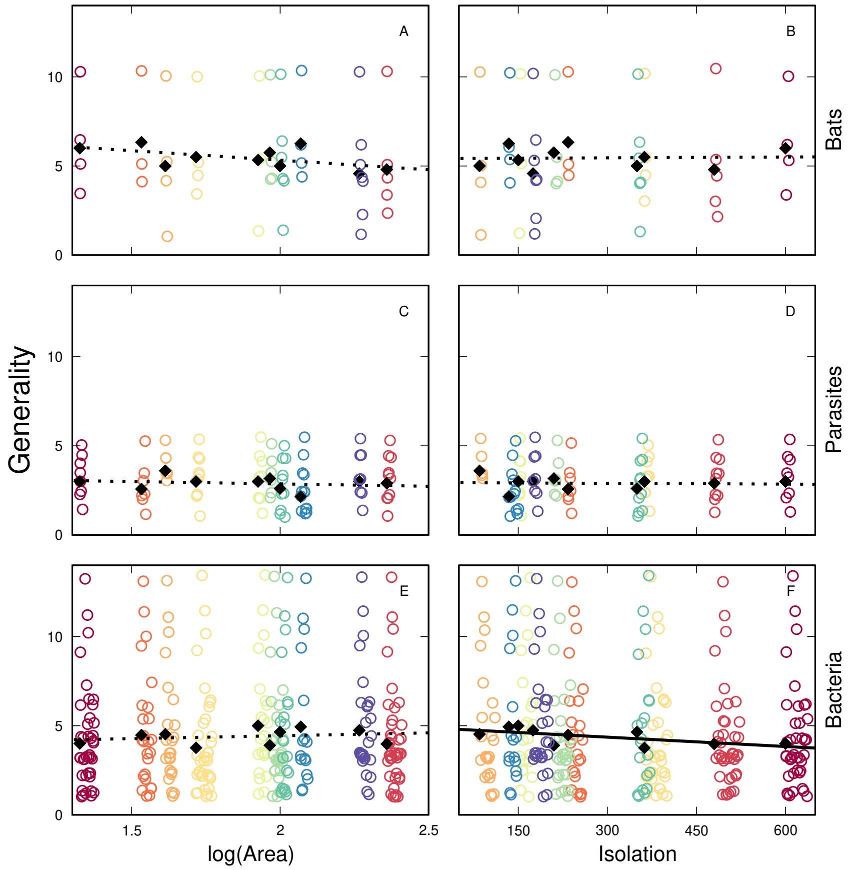


**Figure S3:** For bats and bat flies, mean generality of species found in a forest fragment was not significantly related to either fragment size (log(Area)) or isolation. For bacteria, mean generality decreased slightly but significantly with increasing isolation. Lines indicate predictions of linear regressions of mean generality of all fragments against log of area or isolation. Random horizontal and vertical judder has been added to each point to reduce overlap and allow clusters of taxa with similar properties to be clearly seen. Symbol colour indicates forest fragment.

size ($F_{1,8}$=3.82, $p$=0.087, $R^2$=0.323; Fig. S3), isolation ($F_{1,8}$=0.012, $p$=0.915, $R^2$=0.002), or habitat complexity ($F_{3,6}$=0.977, $p$=0.977, $R^2$=0.328). For bat flies there was no relationship between mean generality and log area ($F_{1,8}$=0.421, $p$=0.535, $R^2$=0.050), isolation ($F_{1,8}$=0.030, $p$=0.866, $R^2$=0.004), or habitat complexity ($F_{3,6}$=0.106, $p$=0.953, $R^2$=0.050). For bacteria there was no relationship between mean generality and fragment size ($F_{1,8}$=0.458, $p$=0.518, $R^2$=0.054) or habitat complexity ($F_{3,6}$=1.00, $p$=0.455, $R^2$=0.333). However, mean generality of bacteria was lower in more-isolated fragments

($F_{1,8}$=5.51, $p$=0.047, $R^2$=0.408).

### Appendix 3: Redundancy and fragment properties

#### Methods

To test for loss of redundancy, we first calculated the mean number of shared partners in each trophic group as an estimate of redundancy using the 'networklevel' function in the R (R Core Team, 2023) package *bipartite* (Dormann *et al.*, 2008). If, on average, more taxa share the same interaction partners, then there is more redundancy in the network. After calculating redundancy for each trophic level in each observed network, we then compared these values to 1000 randomisations of each network (created using the function 'nullmodel' from the R (R Core Team, 2023) function *bipartite* (Dormann *et al.*, 2008) with the argument method='r2d'). For each observed network, we conducted four t-tests of whether the observed redundancy of each trophic level was greater or lesser than the mean of simulated redundancies, using the R (R Core Team, 2023) base function 't.test'. As we conduct two tests on each unique mean, we use a confidence level of 0.975 ($\alpha$=0.025) for each test. Next, we tested whether, for each trophic group in each network type, there was a significant relationship between redundancy and log area, isolation, or habitat complexity (described by the first two PCA axes and their interaction). All models were fit using the base R (R Core Team, 2023) function 'lm'.

As an alternative approach, we also examined both binary and weighted nestedness (NODF; wNODF) as a nested network structure where specialists interact with subsets of the partners of generalists can also provide redundancy. We calculated NODF and wNODF for each network using the R (R Core Team, 2023) function 'networklevel' from the *bipartite* (Dormann *et al.*, 2008) package. We then created 1000 randomisations of each network, keeping row and column totals (the number of observed interactions per taxon) constant. Randomised networks were created using the R (R Core Team, 2023) function ‘nullmodel’ with the argument method=“r2d”, also from the *bipartite* (Dormann *et al.*, 2008) package. NODF and wNODF were calculated for each randomised network and used to calculate the significance of nestedness in the observed network (defined as the proporiton of randomised networks with equal or greater nestedness than the observed value).

### Results

#### Significance of observed redundancy

In the herbivory networks, redundancy was significantly lower than expected for both bats and plants at all sites (Table S1). In the parasitism networks, redundancy was significantly lower than expected for both bat flies and bats at all sites (Table S2). In the bacterial networks, redundancy was significantly lower than expected for bacteria and bat flies at most sites, including the metaweb (Table S3). Bacteria did, however, share more hosts at sites F2, F2, and F6 while bat flies shared more bacteria at site F2.

**Table S1:** In the herbivory networks, both bats and plants had significantly fewer shared interaction partners (lower redundancy) than expected based on network randomisations with the same number of observations per taxon.

| | Bats | | | | | Plants | | | | |
|---|---|---|---|---|---|---|---|---|---|---|
| Site | Obs. | $t$>Rand | $p$>Rand | $t$<Rand | $p$<Rand | Obs. | $t$>Rand | $p$>Rand | $t$<Rand | $p$<Rand |
| metaweb | 1.439 | 115.211 | 0.000 | 115.211 | 1.000 | 1.115 | 112.755 | 0.000 | 112.755 | 1.000 |
| REGUA1 | 0.571 | 102.459 | 0.000 | 102.459 | 1.000 | 1.167 | 89.707 | 0.000 | 89.707 | 1.000 |
| REGUA2 | 0.786 | 73.439 | 0.000 | 73.439 | 1.000 | 1.100 | 92.791 | 0.000 | 92.791 | 1.000 |
| REGUA3 | 0.300 | 36.698 | 0.000 | 36.698 | 1.000 | 0.200 | 52.181 | 0.000 | 52.181 | 1.000 |
| F1 | 1.167 | 7.573 | 0.000 | 7.573 | 1.000 | 1.167 | 50.139 | 0.000 | 50.139 | 1.000 |
| F2 | 0.333 | 74.367 | 0.000 | 74.367 | 1.000 | 0.333 | 62.400 | 0.000 | 62.400 | 1.000 |
| F3 | 0.167 | 74.094 | 0.000 | 74.094 | 1.000 | 0.167 | 73.259 | 0.000 | 73.259 | 1.000 |
| F4 | 0.333 | 37.962 | 0.000 | 37.962 | 1.000 | 0.400 | 34.264 | 0.000 | 34.264 | 1.000 |
| F5 | 0.667 | 8.821 | 0.000 | 8.821 | 1.000 | 1.000 | -8.671 | 1.000 | -8.671 | 0.000 |
| F6 | 1.000 | 40.889 | 0.000 | 40.889 | 1.000 | 0.500 | 82.078 | 0.000 | 82.078 | 1.000 |
| F7 | 0.733 | 92.840 | 0.000 | 92.840 | 1.000 | 0.607 | 113.679 | 0.000 | 113.679 | 1.000 |
| F8 | 1.000 | 49.377 | 0.000 | 49.377 | 1.000 | 0.600 | 72.456 | 0.000 | 72.456 | 1.000 |
| F9 | 0.905 | 57.965 | 0.000 | 57.965 | 1.000 | 1.000 | 57.183 | 0.000 | 57.183 | 1.000 |
| F10 | 0.600 | 32.643 | 0.000 | 32.643 | 1.000 | 1.000 | 34.592 | 0.000 | 34.592 | 1.000 |

**Table S2:** In the parasitism networks, both bat flies and bats had significantly fewer shared interaction partners (lower redundancy) than expected based on network randomisations with the same number of observations per taxon.

| | Bat flies | | | | | Bats | | | | |
|---|---|---|---|---|---|---|---|---|---|---|
| Site | Obs. | $t$>Rand | $p$>Rand | $t$<Rand | $p$<Rand | Obs. | $t$>Rand | $p$>Rand | $t$<Rand | $p$<Rand |
| metaweb | 0.240 | 544.476 | 0.000 | 544.476 | 1.000 | 0.272 | 505.689 | 0.000 | 505.689 | 1.000 |
| REGUA1 | 0.167 | 226.401 | 0.000 | 226.401 | 1.000 | 0.107 | 225.921 | 0.000 | 225.921 | 1.000 |
| REGUA2 | 0.133 | 222.412 | 0.000 | 222.412 | 1.000 | 0.107 | 225.211 | 0.000 | 225.211 | 1.000 |
| REGUA3 | 0.106 | 228.642 | 0.000 | 228.642 | 1.000 | 0.071 | 229.750 | 0.000 | 229.750 | 1.000 |
| F1 | 0.393 | 154.093 | 0.000 | 154.093 | 1.000 | 0.000 | 165.746 | 0.000 | 165.746 | 1.000 |
| F2 | 0.286 | 155.133 | 0.000 | 155.133 | 1.000 | 0.333 | 148.501 | 0.000 | 148.501 | 1.000 |
| F3 | 0.300 | 134.036 | 0.000 | 134.036 | 1.000 | 0.000 | 126.813 | 0.000 | 126.813 | 1.000 |
| F4 | 0.278 | 195.655 | 0.000 | 195.655 | 1.000 | 0.190 | 207.478 | 0.000 | 207.478 | 1.000 |
| F5 | 0.214 | 123.460 | 0.000 | 123.460 | 1.000 | 0.000 | 155.057 | 0.000 | 155.057 | 1.000 |
| F6 | 0.467 | 72.322 | 0.000 | 72.322 | 1.000 | 0.667 | 63.822 | 0.000 | 63.822 | 1.000 |
| F7 | 0.200 | 247.135 | 0.000 | 247.135 | 1.000 | 0.143 | 247.517 | 0.000 | 247.517 | 1.000 |
| F8 | 0.205 | 332.964 | 0.000 | 332.964 | 1.000 | 0.143 | 333.123 | 0.000 | 333.123 | 1.000 |
| F9 | 0.107 | 170.793 | 0.000 | 170.793 | 1.000 | 0.048 | 161.540 | 0.000 | 161.540 | 1.000 |
| F10 | 0.178 | 243.541 | 0.000 | 243.541 | 1.000 | 0.067 | 190.618 | 0.000 | 190.618 | 1.000 |

**Table S3:** In the microbial networks, both bacteria and bat flies had significantly fewer shared interaction partners (lower redundancy) than expected based on network randomisations with the same number of observations per taxon.

| | Bacteria | | | | | Bat flies | | | | |
|---|---|---|---|---|---|---|---|---|---|---|
| Site | Obs. | $t$>Rand | $p$>Rand | $t$<Rand | $p$<Rand | Obs. | $t$>Rand | $p$>Rand | $t$<Rand | $p$<Rand |
| metaweb | 0.649 | 104.252 | 0.000 | 104.252 | 1.000 | 6.949 | 213.173 | 0.000 | 213.173 | 1.000 |
| REGUA1 | 0.475 | -59.632 | 1.000 | -59.632 | 0.000 | 2.111 | 28.264 | 0.000 | 28.264 | 1.000 |
| REGUA2 | 0.606 | 40.228 | 0.000 | 40.228 | 1.000 | 3.900 | 51.310 | 0.000 | 51.310 | 1.000 |
| REGUA3 | 0.445 | 89.901 | 0.000 | 89.901 | 1.000 | 3.133 | 91.030 | 0.000 | 91.030 | 1.000 |
| F1 | 0.409 | 69.788 | 0.000 | 69.788 | 1.000 | 2.900 | 41.972 | 0.000 | 41.972 | 1.000 |
| F2 | 0.607 | -64.776 | 1.000 | -64.776 | 0.000 | 2.067 | -4.763 | 1.000 | -4.763 | 0.000 |
| F3 | 0.747 | -77.532 | 1.000 | -77.532 | 0.000 | 1.900 | 30.249 | 0.000 | 30.249 | 1.000 |
| F4 | 0.410 | 61.676 | 0.000 | 61.676 | 1.000 | 1.893 | 90.406 | 0.000 | 90.406 | 1.000 |
| F5 | 0.614 | 58.637 | 0.000 | 58.637 | 1.000 | 2.667 | 63.206 | 0.000 | 63.206 | 1.000 |
| F6 | 0.664 | -18.357 | 1.000 | -18.357 | 0.000 | 2.667 | 12.570 | 0.000 | 12.570 | 1.000 |
| F7 | 0.416 | 37.336 | 0.000 | 37.336 | 1.000 | 1.048 | 79.683 | 0.000 | 79.683 | 1.000 |
| F8 | 0.342 | 66.560 | 0.000 | 66.560 | 1.000 | 1.333 | 61.530 | 0.000 | 61.530 | 1.000 |
| F9 | 0.561 | 21.587 | 0.000 | 21.587 | 1.000 | 1.900 | 75.307 | 0.000 | 75.307 | 1.000 |
| F10 | 0.414 | 51.926 | 0.000 | 51.926 | 1.000 | 2.667 | 70.559 | 0.000 | 70.559 | 1.000 |

#### Redundancy and fragment properties

Considering fragment sites only, mean redundancy within a trophic group at a site was not significantly related to the log of fragment area, fragment isolation, or habitat complexity (the first two habitat PCA axes and their interaction) for any trophic level in any network type (Table S4). If the three REGUA sites are included in the same models, bat flies were less redundant in their use of bats in larger fragments ($\beta$=-0.051, $p$=0.024) while bats' redundancy in their parasite communities was related to habitat complexity (Table S5). Redundancy in the herbivory or microbial networks was not related to fragment size, isolation, or habitat complexity.

**Table S4:** Summary statistics for regressions relating redundancy (mean number of shared partners within a group) to fragment properties for each network type and trophic group, excluding the three REGUA control sites. Within each network type, the consumer group shown is first.

| Network | Group | Predictor | $F$ | $p$ | $R^2$ |
|---|---|---|---|---|---|
| Herbivory | Bats | log(Area) | 0.297 | 0.600 | 0.036 |
| Herbivory | Bats | Isolation | 0.685 | 0.432 | 0.079 |
| Herbivory | Bats | Complexity | 0.131 | 0.938 | 0.061 |
| Herbivory | Plants | log(Area) | 0.700 | 0.427 | 0.080 |
| Herbivory | Plants | Isolation | 2.70 | 0.139 | 0.253 |
| Herbivory | Plants | Complexity | 1.27 | 0.366 | 0.388 |
| Parasitism | Bat flies | log(Area) | 5.21 | 0.052 | 0.394 |
| Parasitism | Bat flies | Isolation | 0.295 | 0.602 | 0.036 |
| Parasitism | Bat flies | Complexity | 1.34 | 0.347 | 0.401 |
| Parasitism | Bats | log(Area) | 0.002 | 0.969 | <0.001 |
| Parasitism | Bats | Isolation | 0.178 | 0.684 | 0.022 |
| Parasitism | Bats | Complexity | 15.5 | 0.003 | 0.886 |
| Microbial | Bacteria | log(Area) | 0.288 | 0.606 | 0.035 |
| Microbial | Bacteria | Isolation | 4.42 | 0.069 | 0.356 |
| Microbial | Bacteria | Complexity | 0.443 | 0.731 | 0.181 |
| Microbial | Bat flies | log(Area) | 0.304 | 0.597 | 0.037 |
| Microbial | Bat flies | Isolation | 1.25 | 0.296 | 0.135 |
| Microbial | Bat flies | Complexity | 0.485 | 0.705 | 0.195 |

**Table S5:** Summary statistics for regressions relating redundancy (mean number of shared partners within a group) to site properties (including the REGUA control sites) for each network type and trophic group. Within each network type, the consumer group shown is first.

| Network | Group | Predictor | $F$ | $p$ | $R^2$ |
|---|---|---|---|---|---|
| Herbivory | Bats | log(Area) | 0.259 | 0.621 | 0.023 |
| Herbivory | Bats | Isolation | 1.29 | 0.279 | 0.105 |
| Herbivory | Bats | Complexity | 0.236 | 0.869 | 0.073 |
| Herbivory | Plants | log(Area) | 0.529 | 0.482 | 0.046 |
| Herbivory | Plants | Isolation | 0.654 | 0.436 | 0.056 |
| Herbivory | Plants | Complexity | 1.12 | 0.393 | 0.271 |
| Parasitism | Bat flies | log(Area) | 6.82 | 0.024 | 0.383 |
| Parasitism | Bat flies | Isolation | 2.34 | 0.155 | 0.175 |
| Parasitism | Bat flies | Complexity | 3.69 | 0.056 | 0.551 |
| Parasitism | Bats | log(Area) | 0.244 | 0.631 | 0.022 |
| Parasitism | Bats | Isolation | 0.014 | 0.907 | 0.001 |
| Parasitism | Bats | Complexity | 14.9 | $<$0.001 | 0.832 |
| Microbial | Bacteria | log(Area) | 0.058 | 0.814 | 0.005 |
| Microbial | Bacteria | Isolation | 2.91 | 0.116 | 0.209 |
| Microbial | Bacteria | Complexity | 0.291 | 0.831 | 0.089 |
| Microbial | Bat flies | log(Area) | 3.61 | 0.084 | 0.247 |
| Microbial | Bat flies | Isolation | 0.081 | 0.781 | 0.007 |
| Microbial | Bat flies | Complexity | 1.34 | 0.323 | 0.308 |

#### Significance of nestedness

Most of the herbivory networks had significantly higher binary nestedness than randomised versions of the networks (except sites F1 and F5, which had significantly lower binary nestedness than randomised versions of the networks), and all had signficantly greater weighted nestedness than randomisations (Table S6). All of the parasitism networks had significantly higher binary and weighted nestedness than randomised versions of the networks (Table S7). Most of the microbial networks had significantly higher binary nestedness than randomised versions of the networks (Table S8; except sites REGUA1, F2, and F3, which had significantly lower binary nestedness than randomised versions of the networks), and most had significantly higher weighted nestedness than randomised versions of the networks (except sites REGUA2, F2, F3, F6, and F10, which had significantly lower weighted nestedness than randomised versions of the networks).

**Table S6:** Observed binary (NODF) and weighted (wNODF) nestedness values for each parasitism network, as well as test statistics and $p$-values for $t$-tests comparing the observed values to the NODF and wNODF of 1000 randomisations of each observed network.

| | Binary Nestedness | | | | | Weighted Nestedness | | | | |
|---|---|---|---|---|---|---|---|---|---|---|
| Site | Obs. | $t>$Rand | $p>$Rand | $t<$Rand | $p<$Rand | Obs. | $t>$Rand | $p>$Rand | $t<$Rand | $p<$Rand |
| metaweb | 58.8 | 94.2 | 0.000 | 94.2 | 1.000 | 44.3 | 92.8 | 0.000 | 92.8 | 1.000 |
| REGUA1 | 27.8 | 126.8 | 0.000 | 126.8 | 1.000 | 16.7 | 69.6 | 0.000 | 69.6 | 1.000 |
| REGUA2 | 53.5 | 55.4 | 0.000 | 55.4 | 1.000 | 30.3 | 70.8 | 0.000 | 70.8 | 1.000 |
| REGUA3 | 15.0 | 54.6 | 0.000 | 54.6 | 1.000 | 10.0 | 33.3 | 0.000 | 33.3 | 1.000 |
| F1 | 83.3 | -31.2 | 1.000 | -31.2 | 0.000 | 29.2 | 50.3 | 0.000 | 50.3 | 1.000 |
| F2 | 33.3 | 66.4 | 0.000 | 66.4 | 1.000 | 0.0 | 87.5 | 0.000 | 87.5 | 1.000 |
| F3 | 16.7 | 71.1 | 0.000 | 71.1 | 1.000 | 8.3 | 65.2 | 0.000 | 65.2 | 1.000 |
| F4 | 31.3 | 25.8 | 0.000 | 25.8 | 1.000 | 12.5 | 32.4 | 0.000 | 32.4 | 1.000 |
| F5 | 66.7 | -8.7 | 1.000 | -8.7 | 0.000 | 0.0 | 61.5 | 0.000 | 61.5 | 1.000 |
| F6 | 50.0 | 35.3 | 0.000 | 35.3 | 1.000 | 31.3 | 19.4 | 0.000 | 19.4 | 1.000 |
| F7 | 35.3 | 102.6 | 0.000 | 102.6 | 1.000 | 19.4 | 98.9 | 0.000 | 98.9 | 1.000 |
| F8 | 37.5 | 84.7 | 0.000 | 84.7 | 1.000 | 21.9 | 72.1 | 0.000 | 72.1 | 1.000 |
| F9 | 50.9 | 56.4 | 0.000 | 56.4 | 1.000 | 16.4 | 94.2 | 0.000 | 94.2 | 1.000 |
| F10 | 46.2 | 23.1 | 0.000 | 23.1 | 1.000 | 15.4 | 132.2 | 0.000 | 132.2 | 1.000 |

**Table S7:** Observed binary (NODF) and weighted (wNODF) nestedness values for each parasitism network, as well as test statistics and $p$-values for $t$-tests comparing the observed values to the NODF and wNODF of 1000 randomisations of each observed network.

| | Binary Nestedness | | | | | Weighted Nestedness | | | | |
|---|---|---|---|---|---|---|---|---|---|---|
| Site | Obs. | $t>$Rand | $p>$Rand | $t<$Rand | $p<$Rand | Obs. | $t>$Rand | $p>$Rand | $t<$Rand | $p<$Rand |
| metaweb | 11.6 | 615.0 | 0.000 | 615.0 | 1.000 | 5.1 | 458.8 | 0.000 | 458.8 | 1.000 |
| REGUA1 | 7.0 | 220.8 | 0.000 | 220.8 | 1.000 | 5.0 | 110.5 | 0.000 | 110.5 | 1.000 |
| REGUA2 | 5.5 | 206.5 | 0.000 | 206.5 | 1.000 | 1.4 | 148.7 | 0.000 | 148.7 | 1.000 |
| REGUA3 | 3.7 | 241.2 | 0.000 | 241.2 | 1.000 | 2.7 | 147.4 | 0.000 | 147.4 | 1.000 |
| F1 | 0.0 | 278.8 | 0.000 | 278.8 | 1.000 | 0.0 | 123.6 | 0.000 | 123.6 | 1.000 |
| F2 | 0.0 | 201.4 | 0.000 | 201.4 | 1.000 | 0.0 | 140.4 | 0.000 | 140.4 | 1.000 |
| F3 | 0.0 | 163.2 | 0.000 | 163.2 | 1.000 | 0.0 | 81.4 | 0.000 | 81.4 | 1.000 |
| F4 | 10.8 | 190.8 | 0.000 | 190.8 | 1.000 | 3.5 | 149.6 | 0.000 | 149.6 | 1.000 |
| F5 | 0.0 | 183.8 | 0.000 | 183.8 | 1.000 | 0.0 | 115.4 | 0.000 | 115.4 | 1.000 |
| F6 | 22.2 | 96.8 | 0.000 | 96.8 | 1.000 | 16.7 | 59.6 | 0.000 | 59.6 | 1.000 |
| F7 | 0.0 | 370.5 | 0.000 | 370.5 | 1.000 | 0.0 | 261.1 | 0.000 | 261.1 | 1.000 |
| F8 | 10.9 | 337.0 | 0.000 | 337.0 | 1.000 | 5.9 | 226.1 | 0.000 | 226.1 | 1.000 |
| F9 | 6.1 | 181.7 | 0.000 | 181.7 | 1.000 | 2.0 | 151.2 | 0.000 | 151.2 | 1.000 |
| F10 | 7.5 | 254.4 | 0.000 | 254.4 | 1.000 | 5.8 | 153.1 | 0.000 | 153.1 | 1.000 |

**Table S8:** Observed binary (NODF) and weighted (wNODF) nestedness values for each microbial network, as well as test statistics and $p$-values for $t$-tests comparing the observed values to the NODF and wNODF of 1000 randomisations of each observed network.

| | Binary Nestedness | | | | | Weighted Nestedness | | | | |
|---|---|---|---|---|---|---|---|---|---|---|
| Site | Obs. | $t>$Rand | $p>$Rand | $t<$Rand | $p<$Rand | Obs. | $t>$Rand | $p>$Rand | $t<$Rand | $p<$Rand |
| metaweb | 34.6 | 77.182 | 0.000 | 77.182 | 1.000 | 14.0 | 89.799 | 0.000 | 89.799 | 1.000 |
| REGUA1 | 24.8 | -18.299 | 1.000 | -18.299 | 0.000 | 12.0 | 18.819 | 0.000 | 18.819 | 1.000 |
| REGUA2 | 35.6 | 48.065 | 0.000 | 48.065 | 1.000 | 22.2 | -12.935 | 1.000 | -12.935 | 0.000 |
| REGUA3 | 26.2 | 84.737 | 0.000 | 84.737 | 1.000 | 12.6 | 117.113 | 0.000 | 117.113 | 1.000 |
| F1 | 22.6 | 85.639 | 0.000 | 85.639 | 1.000 | 10.0 | 23.851 | 0.000 | 23.851 | 1.000 |
| F2 | 34.9 | -37.833 | 1.000 | -37.833 | 0.000 | 18.0 | -35.566 | 1.000 | -35.566 | 0.000 |
| F3 | 34.0 | -13.274 | 1.000 | -13.274 | 0.000 | 21.0 | -45.849 | 1.000 | -45.849 | 0.000 |
| F4 | 25.5 | 61.221 | 0.000 | 61.221 | 1.000 | 14.2 | 37.872 | 0.000 | 37.872 | 1.000 |
| F5 | 33.6 | 86.969 | 0.000 | 86.969 | 1.000 | 26.0 | 37.592 | 0.000 | 37.592 | 1.000 |
| F6 | 23.1 | 8.465 | 0.000 | 8.465 | 1.000 | 16.5 | -22.244 | 1.000 | -22.244 | 0.000 |
| F7 | 25.6 | 45.495 | 0.000 | 45.495 | 1.000 | 14.0 | 8.708 | 0.000 | 8.708 | 1.000 |
| F8 | 24.2 | 63.727 | 0.000 | 63.727 | 1.000 | 17.7 | 7.107 | 0.000 | 7.107 | 1.000 |
| F9 | 25.1 | 92.016 | 0.000 | 92.016 | 1.000 | 17.3 | 15.246 | 0.000 | 15.246 | 1.000 |
| F10 | 23.7 | 77.142 | 0.000 | 77.142 | 1.000 | 13.1 | -21.662 | 1.000 | -21.662 | 0.000 |

#### Nestedness and fragment properties

When excluding the three REGUA sites, neither binary nor weighted NODF was related to fragment area, isolation, or habitat complexity for any network type (Table S9). When the three REGUA sites were included, binary and weighted nestedness in the parasitism networks was related to habitat complexity (Table S10).

**Table S9:** Summary statistics for regressions relating binary and weighted nestedness (NODF and wNODF) to fragment properties for each network type and trophic group, excluding the three REGUA control sites. Within each network type, the consumer group shown is first.

| Network | Predictor | $F$ | $p$ | $R^2$ | $F$ | $p$ | $R^2$ |
|---|---|---|---|---|---|---|---|
| Herbivory | log(Area) | 0.0838 | 0.780 | 0.010 | 0.0639 | 0.807 | 0.008 |
| Herbivory | Isolation | 2.56 | 0.148 | 0.243 | 1.50 | 0.255 | 0.158 |
| Herbivory | Complexity | 0.604 | 0.636 | 0.232 | 1.64 | 0.277 | 0.451 |
| Parasitism | log(Area) | 1.36 | 0.277 | 0.145 | 1.23 | 0.300 | 0.133 |
| Parasitism | Isolation | 0.110 | 0.749 | 0.0136 | 0.0675 | 0.802 | 0.00836 |
| Parasitism | Complexity | 4.30 | 0.0611 | 0.682 | 4.05 | 0.0684 | 0.670 |
| Microbial | log(Area) | 1.06 | 0.334 | 0.116 | 0.0795 | 0.785 | 0.00984 |
| Microbial | Isolation | 3.42 | 0.102 | 0.300 | 18.3 | 0.00269 | 0.696 |
| Microbial | Complexity | 0.941 | 0.478 | 0.320 | 0.709 | 0.581 | 0.262 |

**Table S10:** Summary statistics for regressions relating binary and weighted nestedness (NODF and wNODF) to fragment properties for each network type and trophic group, including the three REGUA control sites. Within each network type, the consumer group shown is first.

| Network | Predictor | $F$ | $p$ | $R^2$ | $F$ | $p$ | $R^2$ |
|---|---|---|---|---|---|---|---|
| Herbivory | log(Area) | 1.16 | 0.305 | 0.0953 | 0.306 | 0.591 | 0.027 |
| Herbivory | Isolation | 4.01 | 0.0706 | 0.267 | 0.531 | 0.481 | 0.0461 |
| Herbivory | Complexity | 1.03 | 0.426 | 0.255 | 1.90 | 0.201 | 0.387 |
| Parasitism | log(Area) | 0.0368 | 0.851 | 0.00333 | 0.0182 | 0.895 | 0.00166 |
| Parasitism | Isolation | 0.0758 | 0.788 | 0.00684 | 0.0323 | 0.861 | 0.00292 |
| Parasitism | Complexity | 5.24 | 0.023 | 0.636 | 5.17 | 0.024 | 0.633 |
| Microbial | log(Area) | 0.0719 | 0.794 | 0.00649 | 0.101 | 0.757 | 0.00905 |
| Microbial | Isolation | 3.19 | 0.102 | 0.225 | 4.23 | 0.0642 | 0.278 |
| Microbial | Complexity | 0.828 | 0.511 | 0.216 | 0.641 | 0.608 | 0.176 |

### Appendix 4: Summary of samples

**Table S11:** Number of bat individuals collected per species.

| Species | No. captures |
|---|---|
| *Artibeus fimbriatus* | 3 |
| *Artibeus lituratus* | 74 |
| *Artibeus obscurus* | 17 |
| *Carollia perspicillata* | 194 |
| *Chiroderma doriae* | 1 |
| *Dermanura cinerea* | 3 |
| *Glossophaga soricina* | 2 |
| *Platyrrhinus lineatus* | 14 |
| *Pygoderma bilabiatum* | 1 |
| *Sturnira lilium* | 29 |
| *Sturnira tildae* | 3 |
| *Vampyressa pussila* | 13 |

**Table S12:** Number of bat fly individuals collected from sampled bats, per species.

| Species | No. collected | Species | No. collected |
|---|---|---|---|
| *Anastrebla modestini* | 14 | *Strebla guajiro* | 50 |
| *Aspidoptera falcata* | 63 | *Strebla mirabilis* | 1 |
| *Aspidoptera phyllostomatus* | 15 | *Strebla wiedemanni* | 27 |
| *Basilia juquiensis* | 25 | *Trichobius angulatus* | 2 |
| *Exastinion clovisi* | 32 | *Trichobius dugesioides* | 57 |
| *Megistopoda aranea* | 2 | *Trichobius furmani* | 4 |
| *Megistopoda proxima* | 38 | *Trichobius joblingi* | 303 |
| *Metalasmus pseudopterus* | 1 | *Trichobius longipes* | 44 |
| *Neotrichobius delicatus* | 9 | *Trichobius* sp. 1 | 2 |
| *Paraeuctenodes similis* | 5 | *Trichobius* sp. 2 | 5 |
| *Paratrichobius longicrus* | 82 | *Trichobius* sp. 3 | 12 |
| *Paratrichobius* sp. | 5 | *Trichobius* sp. 4 | 5 |
| *Speiseria ambigua* | 39 | | |

### Appendix 5: Species richness and fragment properties

The best-fit model for bat species richness included only an intercept (Table S13). Models including log area, habitat axis 1, or habitat axis 2 had $\delta AIC_c$<2 from the intercept-only model. Bat richness was higher in larger fragments and positively correlated with habitat axis 2 but negatively correlated with habitat axis 1. Parasite richness was best predicted by bat richness, though models including log area or habitat axis 1 were within $\delta AIC_c$<2 of the best-fit model. Parasite richness was higher in fragments with greater bat richness and in larger fragments but was negatively correlated with habitat axis 1. Bacteria genus richness was best predicted by a model including isolation and log area. No other model was within $\delta AIC_c$<2 of the best-fit model. There were more bacteria genera in larger and more isolated fragments.

**Table S13:** AICc and model parameters for models within δAICc<2 of the best-fit model (model 1) for species richness per forest fragment, determined based on the occurrence matrices. For models including terms other than intercepts, $\beta$ and $p$ refer to the non-intercept term. All terms were centered and scaled prior to model fitting.

| Taxon | Model | Terms | AICc | $\beta$ | p |
|---|---|---|---|---|---|
| Bats | 1 | intercept | 65.7 | 2.37 | <0.001 |
| Bats | 2 | log area | 66.7 | 0.113 | 0.174 |
| Bats | 3 | axis 1 | 66.8 | -0.117 | 0.196 |
| Bats | 4 | axis 2 | 67.6 | 0.087 | 0.332 |
| Parasites | 1 | bats | 62.6 | 0.243 | 0.013 |
| Parasites | 2 | log area | 64.1 | 0.195 | 0.024 |
| Parasites | 3 | axis 1 | 64.2 | -0.212 | 0.033 |
| Bacteria | 1 | isolation, log area | 0.252, 0.368 | 0.252, 0.368 | <0.001, <0.001 |

### Appendix 6: Species and interaction turnover

#### Methods

To assess the variability of species and their interactions between sites, we began by calculating Jaccard similarity (J) in consumer species, potential interactions, and observed interactions between each pair of sites (fragments and REGUA sites). Jaccard similarity is simply the number of species common to both fragments divided by the total number of species found at either, or both, sites. Potential interactions were defined as any interaction occurring in the metaweb where the two species involved were observed at the focal site. For herbivory interactions, we assume all plant genera were accessible to bats at any site as plants were not directly sampled. For parasitism and microbial interactions, we considered taxa to be present only if they were observed in this study. Observed interactions were those included in the interaction networks described in the main text. Note that, because not all samples yielded interaction data, some potential interactions may have occurred but not been observed and our measure of species turnover based on all samples may differ slightly from species turnover calculated from the interaction networks.

To uncover whether changes in the structure of interaction networks was largely due to changes in community composition, we followed (Poisot *et al.*, 2012). Using the R (R Core Team, 2023) function 'betalink' from the synonymous package (Poisot *et al.*, 2012), we calculated the $\beta$-diversities of species and interactions as well as the proportion of interaction turnover due to rewiring (i.e., not due to species turnover) between each pair of networks of the same type (herbivory, parasitism, or microbial).

### Results

Overall, Jaccard similarity in species turnover between sites was higher for bats and parasites than for microbes. There was no clear difference in similarity between REGUA sites and forest fragments. Similarity in potential interactions between sites was much higher for herbivory interactions than parasitism or microbial interactions because we assume that bats can always access any plant recorded in our dataset. This assumption is likely false but, without data on the plant species found in each fragment, we have no reasonable alternative. Similarity was also higher in potential parasitism interactions than potential microbial interactions, likely because of the higher turnover in the microbial community. Realised interactions were less similar between sites than potential interactions in all cases, but most notably for herbivory interactions due to the assumption of constant plant availability mentioned above.

**Table S14:** Coefficients ($\beta$) and summary statistics for linear models relating the proportion of interaction turnover due to rewiring to the amount of turnover ($\beta$-diversity) in species or interactions.

| Network | $\beta$-diversity | $\beta$ | $F$ | $p$ | $R^2$ |
|---|---|---|---|---|---|
| Herbivory | Species | -0.977 | 23.2 | <0.001 | 0.207 |
| Herbivory | Interaction | 0.339 | 6.65 | 0.012 | 0.070 |
| Parasitism | Species | -0.301 | 4.18 | 0.044 | 0.045 |
| Parasitism | Interaction | 0.332 | 15.9 | <0.001 | 0.152 |
| Bacterial | Species | -0.836 | 27.4 | <0.001 | 0.236 |
| Bacterial | Interaction | 0.010 | 0.011 | 0.916 | <0.001 |

In all three sets of networks, species and interaction turnover was substantially lower between the three REGUA (control) sites than between REGUA and fragment sites or between fragments. In the REGUA sites, differences in interaction were mainly due to interaction rewiring between species that were present in both sites while differences in interactions between control and fragment sites or between fragments could be more variably attributed to changes in species composition or to rewiring.

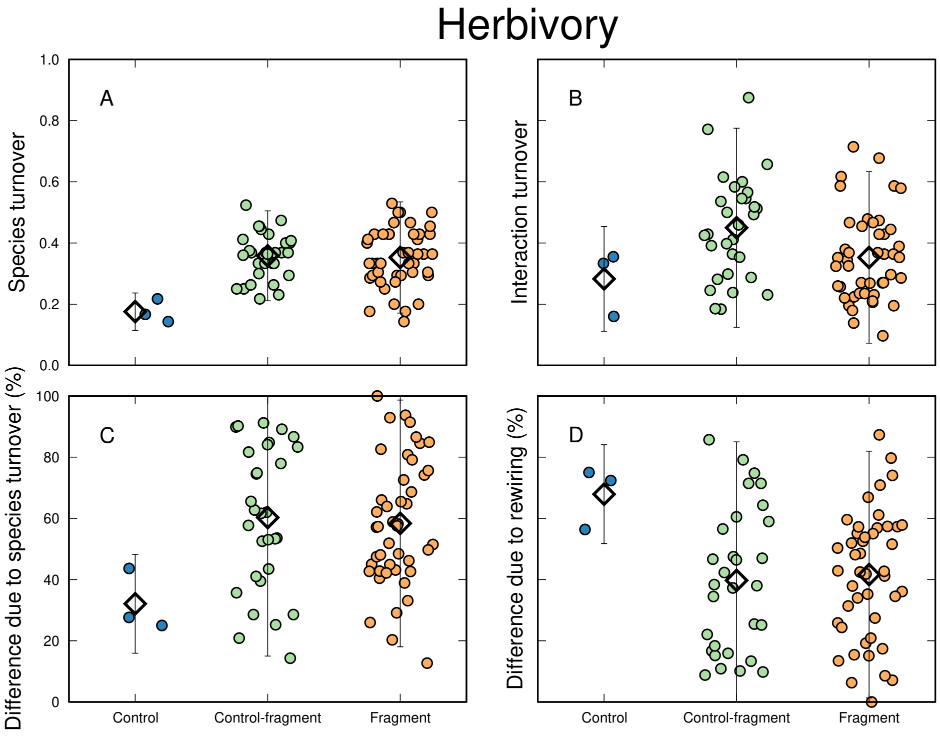


Beta diversity of herbivory networks between sites. We show differences in species composition (A) and interactions (B) as well as the percentage of interaction dissimilarity that can be attributed to changes in species composition (C) or changes in interactions between mutually co-occurring species (D). Black diamonds indicate the mean and error bars indicate a 95% confidence interval.

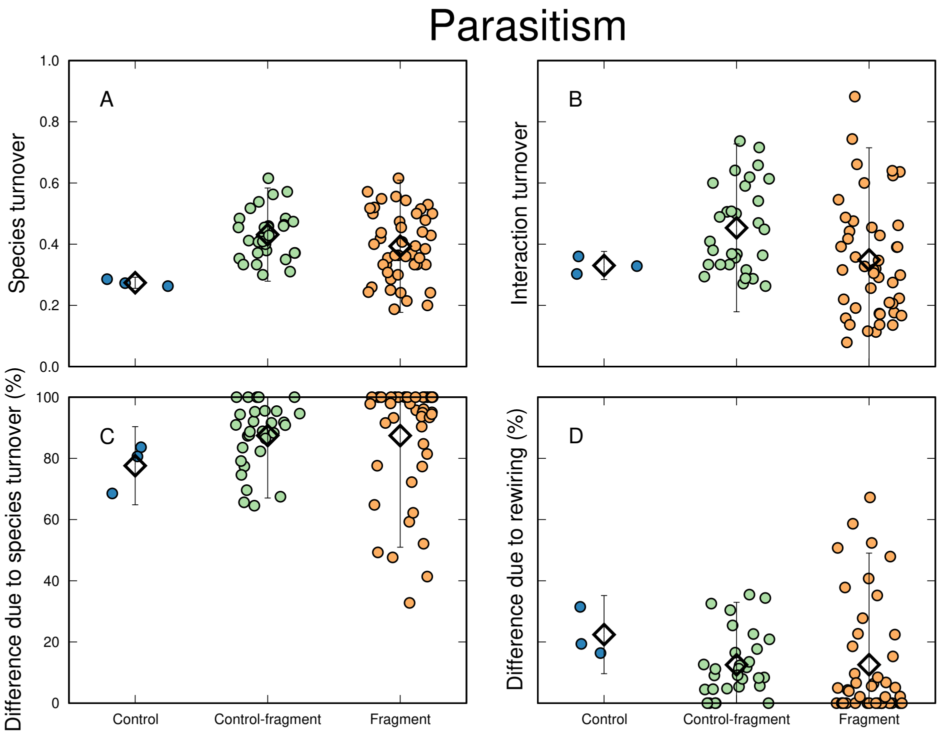


Beta diversity of parasitism networks between sites. We show differences in species composition (A) and interactions (B) as well as the percentage of interaction dissimilarity that can be attributed to changes in species composition (C) or changes in interactions between mutually co-occurring species (D). Black diamonds indicate the mean and error bars indicate a 95% confidence interval.

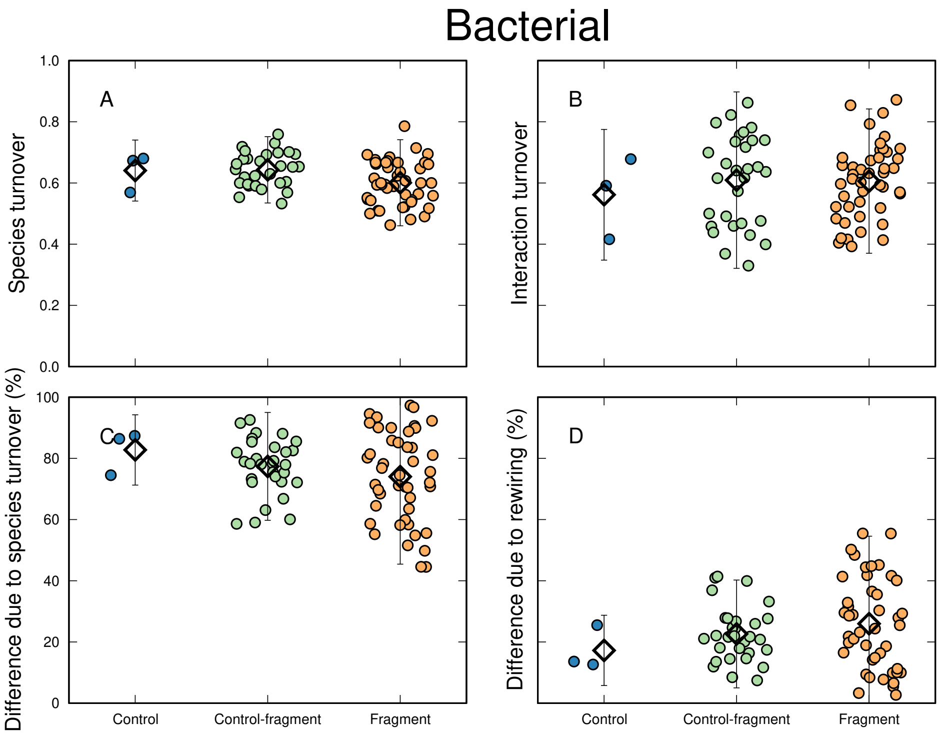


Beta diversity of bacterial networks between sites. We show differences in species composition (A) and interactions (B) as well as the percentage of interaction dissimilarity that can be attributed to changes in species composition (C) or changes in interactions between mutually co-occurring species (D). Black diamonds indicate the mean and error bars indicate a 95% confidence interval.

### Appendix 7: Elements of metacommunity structure (EMS)

#### Methods

For each trophic group (bats, parasites, microbes) we performed an EMS analysis following (Leibold *et al.*, 2004) using the function Metacommunity from the package metacom (Dallas, 2014) for R v. 3.4.1 (R Development Core Team) which uses a hierarchical analysis of a presence/absence matrix of species composition to determine which idealized structure or quasi-structure best fit the data (see Leibold *et al.* (2004); de la Sancha *et al.* (2014) for details). To determine which environmental gradients best represent the primary axes of the EMS analysis we performed a canonical correspondence analysis (CCA; axes shown in Fig. S4) using the function cca in the vegan package for R v3.4.1 (Oksanen *et al.*, 2019). In this analysis the incidence matrix is ordered in the same way as in the EMS analysis, and if coherence is positive and significant a spearman correlation is used to evaluate the relationship between site scores for primary CA axis and the predictor variables. For cases where Clementsian structures are initially identified, we inspected incidence matrices and, where a clear site cluster corresponds to an ecological division, we reconsider sub-groups for independent structures.

#### Results

The metacommunity structure for bats best fit a Clementsian distribution on both axes of ordination (Table S15) which suggests conflicting underlying structures associated with an ecological factor. An inspection of the incidence matrix (Fig. S5) demonstrated a clear division where communities found on the largest fragments cluster with those on continuous forest and have species which are not found in communities found on very small forest fragments (Fig. S5). As a consequence, we refit the EMS to these subsets and found that the bats on small forest fragments best fit a quasi-random species loss model with species loss on the first axis and a quasi-Clementsian model on the second axis, while bats on large fragments best fit a quasi-Glesonian model on both axes. The metacommunity structure for bat flies and microbes also best fit a Clementsian and quasi-Clementsian model, respectively, on both axes though no obvious ecological hypothesis could be used to further split these fragments into groups (Table S15, Fig. S5) thus no further analyses were conducted. CCA found patch area and isolation were strongly associated with the primary axis for bats, parasites and microbes (loadings logArea 0.8839, 0.9768, 0.9976, isolation -0.87, -0.693, -0.545 respectively).

**Table S15:** Elements of Metacommunity Structure results for bats, parasites and microbes in a fragmented landscape of lowland Atlantic Forest in the Guapiaçú River Basin of southeast Brazil. $p$-values in bold are significant ($\alpha$=0.05).

| | | Coherence | | | Species Turnover | | | Boundary Clumping | | |
|---|---|---|---|---|---|---|---|---|---|---|
| Species | Axis | Abs | $p$ | Mean | Rep | $p$ | Mean | I | $p$ | Structure |
| All Bats | 1 | 82 | **<0.001** | 157.48 | 764 | **0.005** | 476.99 | 1.57 | **0.015** | Clementsian |
| | 2 | 105 | **<0.001** | 156.96 | 647 | **0.006** | 375.62 | 1.70 | **0.005** | Clementsian |
| Bats - small frag. | 1 | 18 | **<0.001** | 35.10 | 55 | 0.888 | 58.35 | 1.21 | 0.152 | Quasi-random loss |
| | 2 | 18 | **<0.001** | 37.43 | 32 | 0.758 | 27.33 | 2.22 | **0.008** | Quasi-Clementsian |
| Bats - large frag. | 1 | 25 | **<0.001** | 68.91 | 277 | 0.130 | 220.28 | 1.06 | 0.228 | Quasi-Gleasonian |
| | 2 | 24 | **<0.001** | 69.79 | 173 | 0.140 | 137.35 | 1.20 | 0.092 | Quasi-Gleasonian |
| Parasites | 1 | 68 | **<0.001** | 157.17 | 854 | **0.048** | 643.43 | 1.49 | **0.008** | Clementsian |
| | 2 | 100 | **<0.001** | 154.58 | 754 | **0.006** | 463.01 | 1.68 | **0.011** | Clementsian |
| Microbes | 1 | 868 | **<0.001** | 1564.37 | 31953 | 0.093 | 28669.34 | 1.31 | **<0.001** | Quasi-Clementsian |
| | 2 | 903 | **<0.001** | 1566.26 | 31332 | 0.225 | 29305.02 | 1.29 | **<0.001** | Quasi-Clementsian |

### Discussion

The metacommunity framework assumes that species experience their environment on the same spatial scale and only the abiotic environment determines the suitability of a patch (Guzman *et al.*, 2019). This is at odds with the reality of multi-trophic communities where existence is determined by both abiotic and biotic co-existence and the metacommunity structure at one level may not easily be explained without knowledge of the others (Guzman *et al.*, 2019). In our case, bat flies interact with their abiotic environment (e.g., by depositing eggs on a roosting substrate, thermal tolerance) but can only survive in an environment that also contains a suitable bat host (bottom-up control) and may also experience top-down regulatory control from associated bacteria (e.g., through nutrient provisioning or pathogen invasion). As a consequence, environmental suitability may vary in space and time even if the underlying habitat does not change. Bat-bat fly and bat fly-bacteria communities may follow a consumer resource model but also exist in a hierarchy of spatial processes from landscape through the complexity of host interactions and to the host itself. These hierarchical interactions are understudied within the context of habitat loss and fragmentation.

Metacommunity theory considers a narrow range of dispersal dynamics when, in reality, movement is much more complex and may be tied to life stages, foraging, periodic movements, migration or commuting between patches. In an analysis of host-parasitoid relationships (Holt & Hoopes, 2005), it was demonstrated that differences in dispersal have huge impacts on metacommunity. When evaluating metacommunity it is important to consider the wide variety of habitat and community characteristics which may contribute to the underlying structure and dynamics of the system. An excellent review (Brown *et al.*, 2017) discussed problems associated with too narrowly focussing on the contributions of classic paradigms of species sorting, neutral dynamics, patch dynamics and mass effects to community assembly. Although investigating the processes that create the metacommunity structure is beyond the scope of our analysis, we might suspect that the large reserve areas of REGUA are a source, and small patches a sink leading to classic source-sink dynamics and, at least within the bats, there is some evidence that this is mediated by the habitat structure of small patches. Even in variable sized patches we captured several species that are considered sensitive to habitat fragmentation, including *Trachops cirrhosus*, *Peropteryx macrotis* and *Histiotus velatus* (Emmons & Feer, 1997; Reis *et al.*, 2013). These findings imply that even small forest fragments may be valuable if they provide the specific habitat requirements to support species of special conservation concern. On the other hand, our data also provide clear evidence that a wide variety of highly complex interactions both with other species and the environment, often tied to life stage, may contribute to community structure far beyond the classic pillars of metacommunity theory (Brown *et al.*, 2017).

Our data suggest that all three metacommunities best fit a Clementsian structure with distinct communities replaced with highly coincident ranges. However, there is considerable species-level variation reflecting different environmental tolerances in all three communities. Among bats there is an obvious division between “large” fragments, where communities are more similar to pristine habitat and show a Gleasonian structure, with maximised individualised species’ responses, and “small” fragments structured by random species loss. This suggests the smaller fragments have lost ecological structure associated with more intact habitat. One potential explanation is species specific tolerances to disturbance and dispersal ability between forest patches (Kadmon, 1995; Esbérard *et al.*, 2017). In a complex of islands inside an artificial lake in Panama small island bat communities represent a subset of the communities in bigger islands and mainland habitat (Meyer & Kalko, 2008). In the Atlantic Forest, while habitat fragments are not as isolated, they may act somewhat like islands for more habitat restricted species. In comparison to the bats, both the bat flies and their associated microbes mirrored the overall Clementsian structure of the bats but lacked any obvious subdivision between large and small fragments. While the incidence matrix for bats (Fig. S5a) clearly shows the communities on the largest fragments are similar in composition to the REGUA sites other communities were less clear in how substructure aligns with ecological factors. We explored various clustering techniques to detect significant Clementsian substructures (analysis not shown) but the outcomes either did not correspond to any obvious ecological hypothesis or simply separated all patches into individual structures. This suggests a more complex ecological substructure and we speculate that it likely involves host identity and phylogenetic history contributing to metacommunity structure along with environmental gradients, but this requires far greater investigation.

One of the most difficult aspects of measuring community assembly is to establish an appropriate scale for analysis (Meynard *et al.*, 2013). For example, some species may migrate or commute over landscapes in a single night, while others travel only very short distances from their natal roost and this may be true within and between trophic levels. Among the bats of the Caribbean islands (Presley & Willig, 2010) metacommunity had a Clementsian structure at the regional-scale, and while bats of the Bahamas and Lesser

Antilles also show a Clementsian structure, bats of the Greater Antilles showed a nested structure (Presley *et al.*, 2010), suggesting that the process of species assembly may vary across different scales within the same taxa. Lack of corresponding subdivisions between our trophic communities suggest these taxa are experiencing the landscape at very different scales. As we hypothesize above, at higher trophic levels, the environmental gradient in our EMS analysis may be a combination of the host itself and abiotic gradients, highlighting the need for a new trophic perspective on metacommunity structure which includes "host" as an element of the ecological gradient. Aside from spatial scale, the use of different taxonomic scales among trophic levels has consequences for overall structure of networks and communities, aside from changing the number of taxa represented. For example, microbial data was generated at ASV level, but genus-level resolution was used throughout microbial networks to avoid mixed taxonomic resolution in the network (Hemprich-Bennett *et al.*, 2021) and due to lack of resolution for microbial data at this locus. Less information can be expressed at higher taxonomic levels as data becomes less diverse, but it is easier to ascertain prominent patterns (Banerjee *et al.*, 2016; Lupatini *et al.*, 2014).

**Figure S4:** A CCA plot for the habitat variables. CCAhab1 explained 34.9% of the variance and showed positive associations with the abundance of grasses, the palm *Astrocaryum* (Irís) and *Cecropia* trees, and a negative association with the abundance of lianas, whereas CCAhab2 explained an additional 26.1% of the variance and was associated with the presence of watercourses, increased overstory and understory, and the presence of fallen logs.

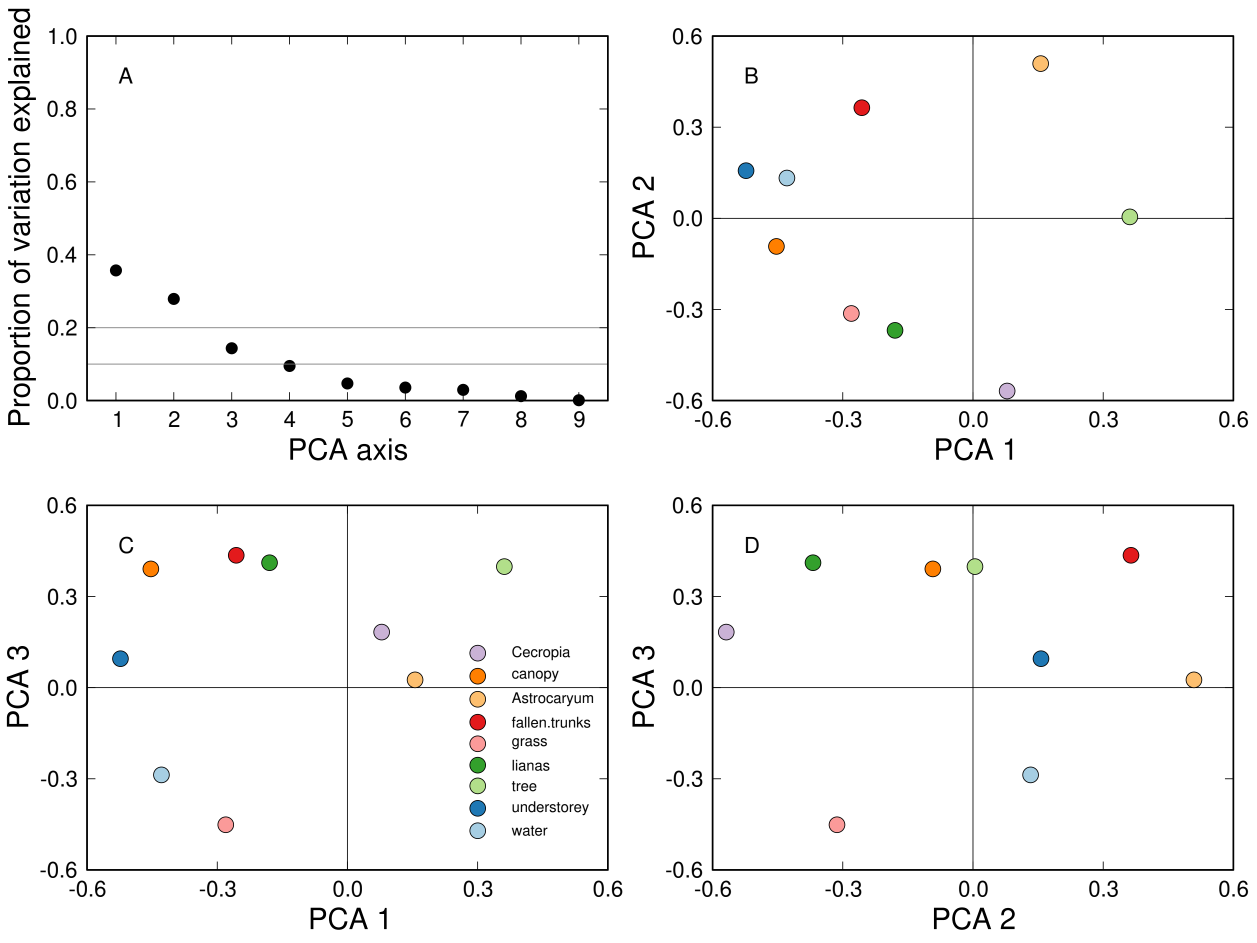

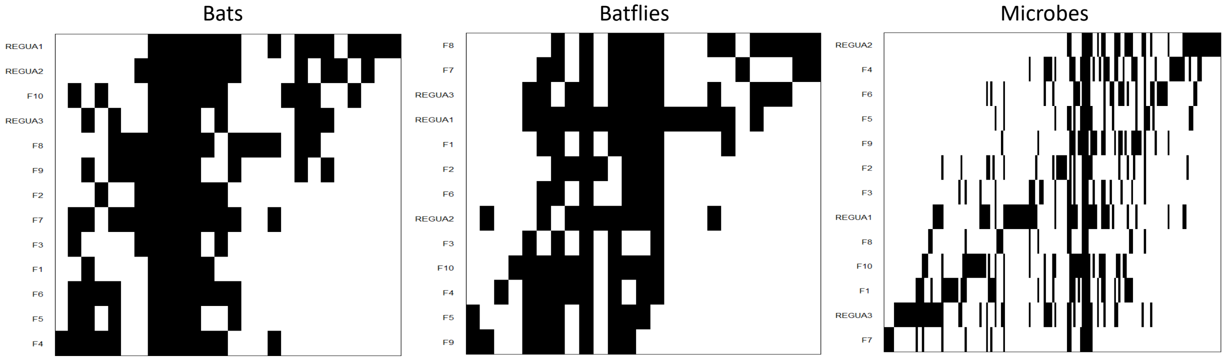


**Figure S5:** Analysis using the elements of metacommunity structure indicates that all three trophic levels of this community have a Clementsian structure. In the case of the bats two groups correspond to the larger sites (F8, F9, F10, REGUA) and the smaller sites (F1-7). Similar ecological hypotheses for clusters are not apparent in the batfly and microbial communities.
